# The obesity-linked human lncRNA AATBC regulates adipocyte plasticity by stimulating mitochondrial dynamics and respiration

**DOI:** 10.1101/2021.08.09.455681

**Authors:** Maude Giroud, Stefan Kotschi, Yun Kwon, Ophélia Le Thuc, Anne Hoffmann, Manuel Gil-Lozano, Michael Karbiener, Juan Carlos Higareda-Almaraz, Sajjad Khani, Daniel Tews, Pamela Fischer-Posovszky, Wenfei Sun, Hua Dong, Adhideb Ghosh, Christian Wolfrum, Martin Wabitsch, Kirsi A. Virtanen, Matthias Blüher, Søren Nielsen, Anja Zeigerer, Cristina García-Cáceres, Marcel Scheideler, Stephan Herzig, Alexander Bartelt

## Abstract

Adipocytes are critical regulators of metabolism and energy balance. While white adipocyte dysfunction is a hallmark of obesity-associated disorders, the activation of thermogenic brown and beige adipocytes is linked to improved cardiometabolic health. As adipocytes dynamically adapt to environmental cues by functionally switching between white and thermogenic phenotypes, a molecular understanding of this adipocyte plasticity could help improving energy balance and weight loss. Here, we show that the long non-coding RNA (lncRNA) Apoptosis associated transcript in bladder cancer (AATBC) is a human-specific regulator of adipocyte plasticity. Searching for new human lncRNAs implicated in adipocyte biology we compared transcriptional profiles of human adipose tissues and cultured adipocytes and discovered that AATBC was enriched in thermogenic conditions. Using primary human adipocytes and immortalized human adipocytes we found that gain-of-function of AATBC enhanced the thermogenic phenotype whereas loss-of-function diminished this effect. The AATBC-mediated increase in mitochondrial respiration was linked to a more fragmented mitochondrial network and vice versa. While we found that AATBC is predominantly located in the nucleus, its effect on global transcription was only marginal. As AATBC is specific to humans, we expressed AATBC in adipose tissue of mice to study its systemic impact, which led to lower plasma leptin levels. Interestingly, this association was also present in human subjects, as AATBC in adipose tissue was inversely correlated with plasma leptin levels, body mass index and other measures of metabolic health. In conclusion, AATBC is a novel obesity-linked regulator of adipocyte plasticity and mitochondrial function in humans.

## Introduction

Adipocytes are critical regulators of energy metabolism and nutrient homeostasis. As adipocytes are energy reservoirs and secrete endocrine hormonal regulators, they represent an important metabolic buffer system. Therefore, adipocyte dysfunction is tightly linked to metabolic disorders, including obesity, diabetes, and cardiovascular disease. While in humans, metabolic disease is mainly associated with white adipocytes, recent data indicate that thermogenic adipocytes in brown adipose tissue (BAT) play a role in energy homeostasis in humans as well [1–4]. These cells are diverse in nature and display high phenotypic plasticity [5]. Their primary function is non-shivering thermogenesis (NST), mediated by Uncoupling protein 1 (UCP1) and UCP1-independent futile cycling mechanisms in response to several environmental stimuli [6–10].

The development, activity, and maintenance of thermogenic adipocytes are mainly regulated physiologically by cold-induced adrenergic stimulation. When humans are subjected to cold exposure, thermogenic adipocytes increase their metabolic activity, and if cold exposure is sustained, thermogenic adipocytes are also recruited in white adipose tissue (WAT), an adaptive phenomenon referred to as adipose tissue browning [2, 11]. Conversely, when the cold stress is removed thermogenic adipocytes adopt a quiescent phenotype functionally reminiscent of white adipocytes, referred to as adipose tissue whitening [12]. In mice, thermoneutrality is ca. 30 °C [13], but humans are thermoneutral at much lower temperatures and, therefore, humans only display limited thermogenic activity [14], as their warm environment makes NST obsolete. However, in colder regions, people display markedly increased NST [2], indicating that the environmental temperature are major determinant of BAT activity. Furthermore, activating BAT has remarkable benefits for metabolic health in mouse models of metabolic disorders [15–17], but also, in the general human population BAT is linked to better cardiometabolic health [18]. Understanding the molecular switches and hormonal cues regulating thermogenic plasticity in adipocytes, thus dictating the balance of energy storage vs energy dissipation, might help to develop novel therapies for treating obesity-associated metabolic disorders.

Key aspects of thermogenic plasticity include facilitating an increase of mitochondrial respiration in response to cold, and conversely, directing the involution of mitochondrial capacity when the adipocytes return to a non-thermogenic state at thermoneutrality. In most cells, a fragmented mitochondrial network is associated with stress, whereas fused mitochondria are thought to display enhanced respiration [19]. In the case of adipocytes, mitochondrial dynamics are largely regulated by nutrient availability and depend on the thermogenic status of the adipocyte [20–23]. The interplay of molecules governing these mitochondrial adaptations are complex and involve a large panel of transcriptional and posttranslational regulators, for example peroxisome proliferator-activated receptor co-activator 1-alpha (PGC1α) [24], Nuclear factor erythroid-2, like-1 (NFE2L1, also known as NRF1 or TCF11) [25], Mitofusin-1 and -2 (MFN1 and MFN2) [26–28], OPA1 mitochondrial dynamin like GTPase (OPA1) [21, 29] as well as dynamin 1 like (DNM1L, also known as DRP1) [23]. For example, adipocyte-specific Mfn2-deficient mice display blunted mitochondrial fusion and impaired adaptative thermogenesis [27, 28]. In human adipocytes, it has been shown previously that an acute activation of thermogenic adipocytes involves fission of the mitochondrial network for enhanced respiration [23, 26].

In addition to these transcriptional and posttranslational mechanisms, non-coding RNAs and especially long non-coding RNAs (lncRNAs) have emerged as potential regulators of adipocyte function [30–34]. The number of lncRNAs in the genome is correlated with organism complexity, they are highly tissue-specific, and their sequence is only poorly conserved between species [35]. They are found across the entire genome and adopt secondary structures that confer several regulatory properties. lncRNA are longer than 200 nucleotides and bind to many types of molecules including RNA, DNA, amino acids, and proteins [36]. In addition to their interactome, their cellular localization also defines their function. For example, when localized to the nucleus, lncRNA impact transcriptional control, genomic imprinting, and chromatin condensation [37]. Furthermore, in the cytoplasm, lncRNAs are decoys for mRNA translation and miRNA function, thus affecting mRNA turnover [38]. The study of lncRNAs in metabolism is an emerging field. In adipocytes, lncRNA have been linked to adipogenesis [34]. For example, Brown fat lncRNA 1 (Blnc1) and brown adipose tissue–enriched lncRNA 10 (lncBATE10) promote the development of thermogenic adipocytes through the regulation of Early B Cell Factor 2 (EBF2) and PGC1α, respectively [42–44]. CCCTC-binding factor (zinc finger protein)-like, opposite strand (Ctcflos) was described as a regulator of browning via regulation of PR/SET Domain 16 (PRDM16) splicing [45]. However, most of these lncRNAs are specific to mice and are not found in humans. One study found 318 lncRNAs conserved in humans and mice [46]. To the best of our knowledge, LINC00473 is so far the only non-conserved, human lncRNA implicated in the regulation of thermogenic adipocytes[47].

In the current study, we set out to discover new lncRNAs specific to human adipocytes and linked to NST and obesity. Based on a transcriptional comparative analysis between human WAT and BAT as well as adrenergic stimulation of different models of cultured human adipocytes, we show that the lncRNA Apoptosis associated transcript in bladder cancer (AATBC) [48–51], regulates thermogenic plasticity of adipocytes by stimulating mitochondrial dynamics and respiration, and is linked to human obesity.

## Material and Methods

### Cell culture

#### Human SVF (RNAseq screening, figure 1)

Human primary adipose-derived stem cells (hpASCs) were isolated and stimulated as previously described [52]. In brief, subcutaneous lipoaspirates from healthy female donors (n = 4) were thawed, cultivated in EGM-2 Medium (Lonza), and used after 1–3 passages. 2 days after the cells reached confluency (= day 0), adipocyte differentiation was induced with the following medium (DMEM/Ham’s F12 (50:50)) supplemented with 5 mM HEPES, 2 mM L-glutamine, 100 μg/ml normocin, 860 nM insulin, 10 μg/ml apo-transferrin, 100 nM rosiglitazone, 0.2 nM triiodothyronin) supplemented with 100 μM 3-isobutyl-1-methylxanthine (IBMX), and 1 μM dexamethasone (Dex). At day 7 of differentiation, supplementation with insulin was stopped. At day 9 of differentiation, the differentiated adipocytes were stimulated with 1 μM norepinephrine (NE; dissolved in 10 mM HCl) or vehicle (VE, 10 mM HCl) for 3 hours.

#### hMADS cells

Human multipotent adipose derived stem (hMADS) cells were cultured as previously described [53]. Briefly, cells were kept until confluency in proliferation medium (Dulbecco’s Modified Eagle’s Medium (DMEM), (Lonza, Switzerland # BE12-707F) supplemented with 10 % FBS (Sigma, Germany), 10 mmol/l 4-(2-hydroxyethyl)-1-piperazineethanesulfonic acid (HEPES) (Gibco, Germany #15630056), 2.5 ng/ml human fibroblast growth factor-basic (hFGF2) (PeproTech, Germany #100-18B), 50 mg/ml penicillin, and 50 mg/ml streptomycin (Gibco, Germany #15140122), followed by 48 h of incubation without hFGF2. The induction of the differentiation towards adipocytes was started (d0) using a cocktail of DMEM/Ham’s F12 (Lonza, Switzerland # BE12-615F) supplemented with 10 mg/ml apo-transferrin (Sigma, Germany # T1428), 10 nM insulin (Sigma, Germany # I 9278), 100 nM rosiglitazone (Cayman, Germany #71740), 0.2 nM triiodothyronine (T3) (Sigma, Germany #6893-02-3), 1 mmol/l dexamethasone (DEX) (Sigma, Germany # D4902), 1 mM 3-isobutyl-1-methylxanthine (IBMX) (Sigma, Germany #28822-58-4). After 48 h the induction medium was replaced by the differentiation medium containing DMEM/Ham’s F12, 10 mg/ml transferrin, 10 nM insulin, 0.2 nM triiodothyronine and 1 μM rosiglitazone. Until day 9, the medium was changed every other days. At day 9, thermogenic differentiated hMADS cells were obtained with chronic rosiglitazone treatment until day of completion of the experiment (d14).

#### Human SVF purification and differentiation

Human stromal vascular fraction was purified from subcutaneous adipose tissue from patients undergoing abdominoplasty. The study was approved by the ethical committee (vote no. 300/16) of the University of Ulm after written informed consent. Minced tissue was chemically digested using collagenase (Sigma, Germany #11088793001) in BSA (Sigma, Germany #A8806-5G) media (ratio 1:3) for 30-45 min until the digestion was stopped by the addition of FBS (Sigma, Germany) at a 10 % v/v ratio. To purify primary adipocytes from of the digested tissue, the mixture was first filtered consecutively through 100 µm, 70 µm and 48 µm meshes and then centrifugated to finally separate the primary adipocytes in the pellet from the supernatant. Cells were plated and treated like hMADS cells.

#### Human SVF purification and differentiation (refers to figures 2A and 2B)

Preadipocytes isolated from the supraclavicular and abdominal subcutaneous region were cultured, differentiated and stimulated as previously described [54]. RNA was extracted from adipocytes using the Trizol method. RNA sequencing was performed by BGI (Hong Kong) using 1000 ng RNA for the TruSeq cDNA library construction (Illumina). 3Gb data was generated per sample on a HiSeq 2000 sequencer (Illumina) and a 91-paired end sequencing strategy was used. Read quality was assessed using FastQC (http://www.bioinformatics.babraham.ac.uk) and the following pre-processing steps where performed using the Fastx toolkit (http://hannonlab.cshl.edu) and PRINSEQ: 7 nt were clipped off from the 5’-end of every read [55]. The reads were then filtered to remove all N-reads. The 3’
s-ends were then trimmed, and the reads filtered to minimum Q25 and 50 bp length. Reads were then mapped with tophat2 to the human genome GRCh38 Ensembl release 77. Read counts were imported into R, and DESeq2 was used for identifying differential expression.

### Modulation of gene expression

The expression of *AATBC* was modulated in vitro using antisense oligonucleotide-mediated knockdown (ASO, Exiqon, #634048-6, #634048-4), siRNA-mediated knockdown (Lincode Human AATBC (Horizon Discovery, #284837) and adenovirus (AV) mediated overexpression (VectorBuilder, #AVM (VB150925-10024) + GFP). White hMADS were transfected at day 12 of differentiation with 30 nmol of siRNA or ASO at day 10 using Lipofectamine™ RNAiMAX transfection reagent (Thermo Scientific, #13778075) according to the manufacturer’s protocol. For overexpression, white hMADS cells were infected with 200 particles of virus per cell. 24 h later, the transfection medium was removed and replaced by the suitable differentiation medium. 48 h after transfection cells were harvested.

### Nuclear RNA extraction

Nuclear RNA was purified using a Cytoplasmic & Nuclear RNA Purification Kit (Norgen, Canada # 21000) according to the manufacturer’s instructions. cDNA was synthesized with Maxima™ H Master Mix 5x (Thermo Fisher Scientific, #M1661), followed by qPCR according to the PowerUp™ SYBR Green technology (Applied Biosystems, #A25741)

### Immunoblot

Protein expression was investigated using western blot as described previously [53]. Cells were lysed in RIPA buffer (150 mM NaCl, 5 mM EDTA, 50 mM Tris pH 8, 0.1 % w/v SDS, 1 % w/v IGEPAL® CA-630, 0.5 % w/v sodium deoxycholate) containing protease inhibitors (Sigma) in a 1:100 v/v ratio. Protein concentrations of the lysates were determined using Pierce BCA Protein Assay (Thermo Scientific) following the manufacturer’s protocol. 30 µg of total protein were loaded per well on Bolt™ 4–12 % Bis-Tris gels (Thermo Scientific, #NW04120BOX) and blotted on a 0.2 µm PVDF membrane (Bio-Rad, #1704156) using the Trans-Blot® Turbo™ system. Membranes were incubated in primary antibodies at a dilution of 1:1000 v/v in ROTI-Block (Roth, Germany #A151.1) after blocking in ROTI-Block Secondary antibodies (concentration of 1:10000 v/v in ROTI-Block. All antibodies used are shown in Table 1. After antibody incubation membranes were washed 4 x for 10 min in TBS-T (Tween 0.1 %). Bands were detected using a Chemidoc MP Sytstem (Bio-Rad, #1704156).

**Table 1:**
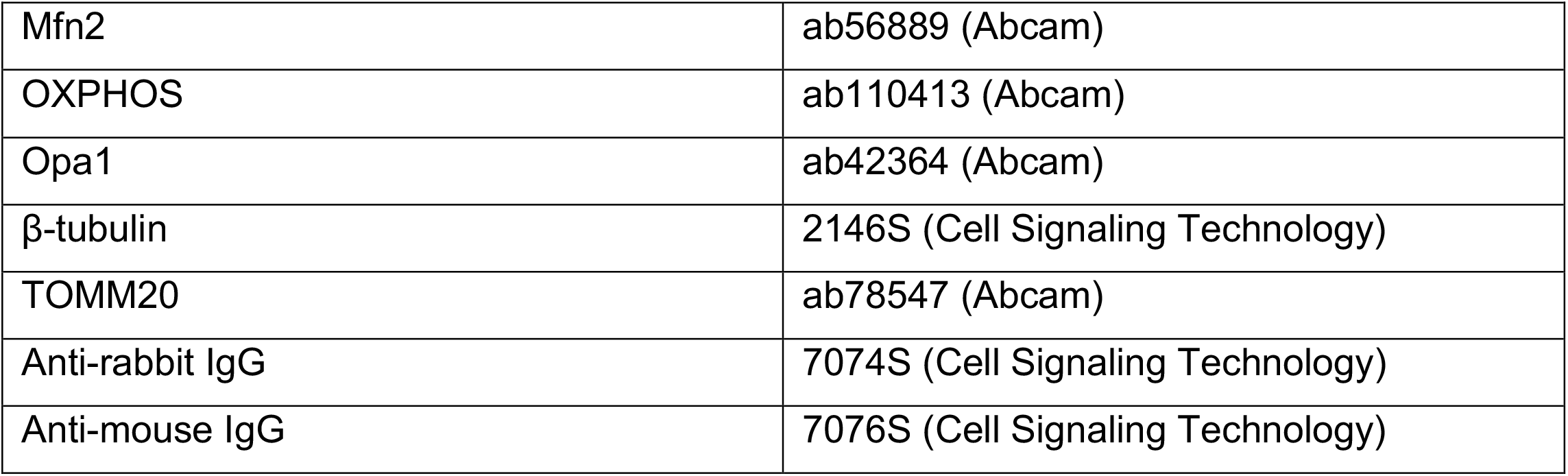
Antibodies:

### Gene expression analysis

RNA of cells and tissues were extracted using TRIzol (Invitrogen, #15596026) and NucleoSpin® RNA kit (Macherey-Nagel, #740955.250) according to the manufacturer’s instruction. cDNA was synthesized with Maxima™ H Master Mix 5 x (Thermo Fisher Scientific, #M1661), followed by qPCR according to the PowerUp™ SYBR Green technology (Applied Biosystems, #A25741) using the primers listed in Table 2. Gene expression was calculated using the ΔΔCt-method normalized to the housekeeping gene indicated in the respective figure.

**Table 2:**
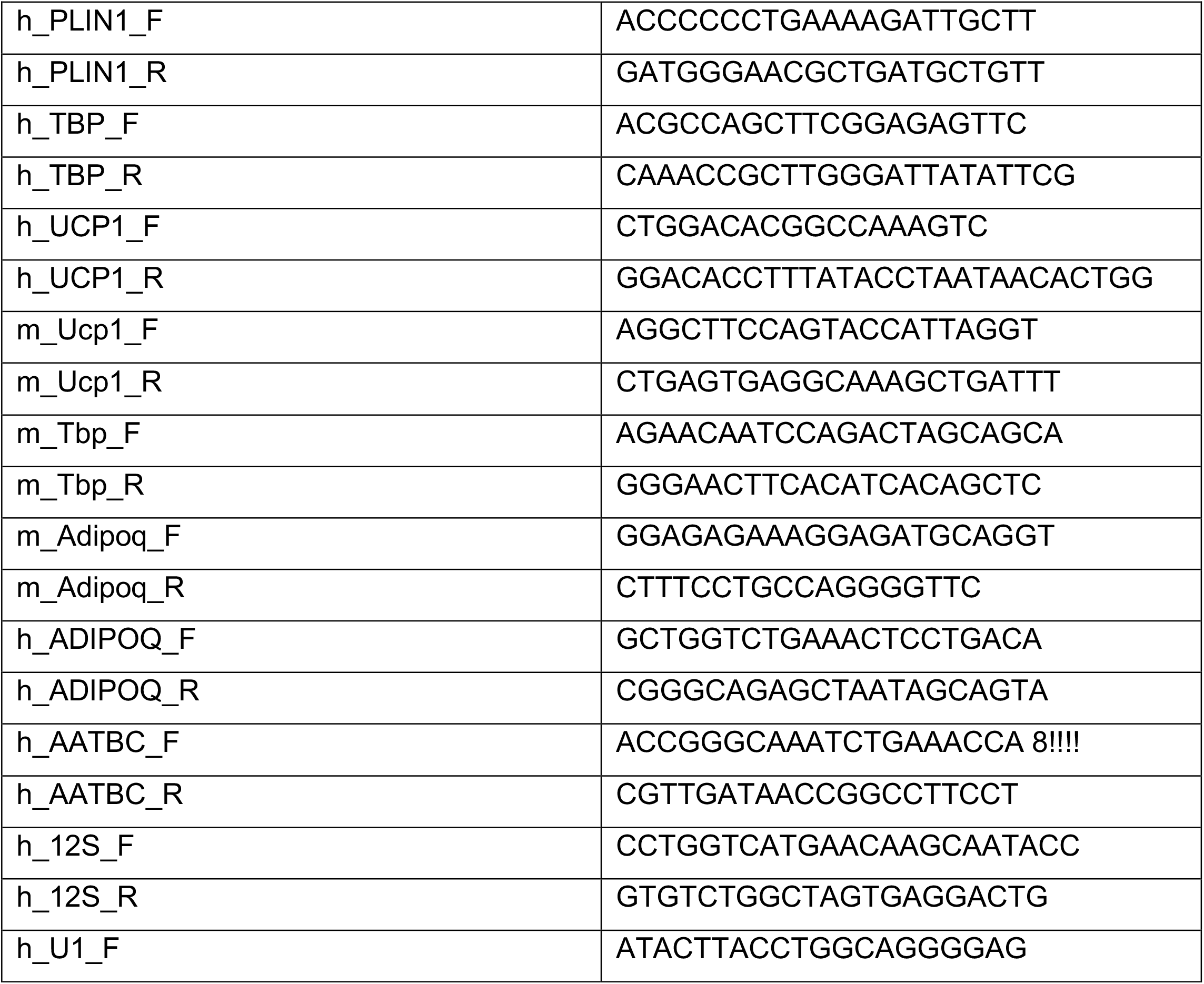
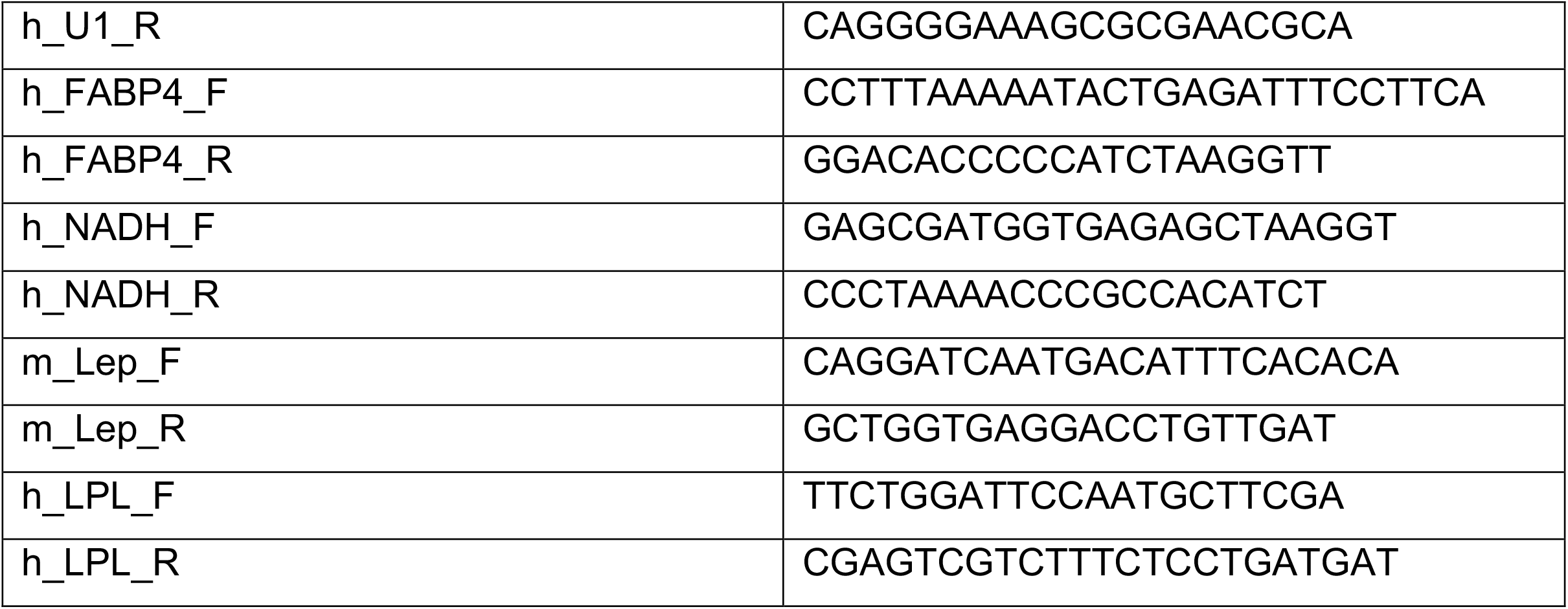
Primers:

### Analysis of mitochondrial respiration

To assess mitochondrial respiration a Seahorse XFe24 device (Agilent # 102238-100, S7801A) was used. hMADS cells were seeded differentiated and transfected or transduced at day 12 as described above. Oxygen consumption rate was determined during a Mito Stress Test (1 µM Oligomycin-A (Sigma-Aldrich, #75351), 1.2 µM FCCP (Sigma-Aldrich, #C2920), 2 µM Rotenone (Sigma-Aldrich, #R8875), 2 µM Antimycin A (Sigma-Aldrich, #A8674)) with additional treatment with Isoproterenol (100 nM, Sigma-Aldrich, #I5627-5G).

### Quantification of mitochondrial DNA

To assess cellular mitochondrial amount, DNA was extracted using a DNA Extraction Kit (Macherey-Nagel, #740952.50) and used for qPCR as described before [56]. The expression of the mitochondrial gene *NADH* was normalized to the single copy nuclear gene *LPL*.

### In situ hybridization - RNAscope®

To visualize the intracellular localization of ATTBC, we used the RNAscope® Multiplex Fluorescent Reagent Kit v2-Hs (ACD Bio, #323135). Undifferentiated hMADS cells were seeded on coverslips in 24-well tissue culture dishes, grown to confluence and differentiated into thermogenic adipocytes for 12 days (see hMADS cells differentiation). *In situ* hybridization was performed according to the manufacturer’s protocol (Advanced Cell Diagnostics), using a probe designed to specifically detect human AATBC (RNAscope® Probe Hs-AATBC (Advanced Cell Diagnostics, #519681)), in combination with the Opal 620 Fluorophore Reagent (Akoya Biosciences). Cells were visualized on a Leica TCS SP5 confocal microscope (Leica, Germany). Images were acquired for all conditions as z-stacks (steps of 0.5 µm along the z axis) with a glycerol-based immersion fluid-immersed 63 x objective. Acquisition parameters were kept consistent for all images. The same image processing was performed using ImageJ (FIJI) software for the different conditions.

### Mitochondria 3D Reconstruction and Morphometric Analysis

Immunofluorescence was performed at day 14 of differentiation as previously described [23]. To visualize the mitochondrial network TOMM20 antibody (Abcam, #ab78547, 1:1000 v/v) was used followed by incubation with a fluorescent antibody (molecular probes, A-21429, 1:1000 v/v). Immunofluorescent samples were analyzed using a Laser Scanning Confocal Microscope (Olympus Fluoview 1200, Olympus, Tokyo, Japan) equipped with an Olympus UPlanSApo 60 × 1.35 and an UplanSApo 40 × 1.25Sil Oil immersion objective (Olympus, Tokyo, Japan) at a resolution of app. 100 µm/pixel (60x) and 600 nm step size. For mitochondrial 3D reconstruction, images were deconvolved using the FIJI plugins point spread function (PSF) generator [57] and DeconvolutionLab [58]. Z-step was set to 0.6 µm and a PSF algorithm (Born & Wolf 3D Optical model) was used for PSF generation, as previously described [59]. The generated PSF and a 3D deconvolution algorithm (Richardson-Lucy with TV regularization) were applied to microscopic images using DeconvolutionLab. From the deconvolved 2D and 3D binary images (8-bit images), mitochondrial network was determined by generating a skeleton of the images using the Fiji plugin Skeletonize3D and analyzed using the plugin AnalyzeSkeleton (2D/3D). This plugin tags all pixel/voxels in a skeleton image and then counts the junctions and branches of the mitochondrial network and measures their average length. For mitochondrial network analysis, minimum of 20 cells were analyzed.

### Mouse experiments

Animal experiments were performed in accordance with German animal welfare legislation and approved by the state ethics committee and government of Upper Bavaria (no. ROB-55.2-2532.Vet_02-17-125). Mice were group-housed at 22 °C with a 12 h dark–light cycle in the animal facility of Helmholtz Center Munich. 9-week-old male C57BL/6J mice were purchased from Janvier Labs and acclimatized in the Helmholtz center facility for 3 weeks. Induction of anesthesia was performed using 4 % isoflurane and upheld by inhalation of 2 % isoflurane. Incisions were made over the intrascapular brown adipose tissue and inguinal adipose tissue. Injections of 25 µl containing 4*10^7^ viral particles of AV_AATBC were performed into each lobe of the tissues. For glucose tolerance test (GTT), performed at day 7 and insulin tolerance test (ITT), performed at day 14, mice were fasted for 6 h after which they received intraperitoneal injections of glucose (3 g/kg for GTT) and insulin (0.7 U/kg for ITT). Blood glucose was measured from the tail vain at 15, 30, 60, 90 and 120 minutes after injection. After 12 days mice were euthanized using cervical dislocation and necropsy was performed. Plasma was analyzed for metabolic parameters including adiponectin, triglycerides and non-esterified fatty acids using a serum analyzer (AU480 Chemistry analyzer Beckman Coulter). Plasma levels of leptin and insulin were assessed using ELISAs used according to the manufacturer’s instructions (R&D Systems, #MOB00B and Crystal Chem, #90082, respectively).

### Human data

#### Human adipose tissue samples (RNAseq screening, refers to figure 1)

Human BAT and WAT were removed from the same incision site in the supraclavicular localization, BAT being fluorodeoxyglucose-positron emission tomography (FDG-PET)-positive scan sites and WAT being not. The Study protocol was approved by the ethics committee of the Hospital District of Southwestern Finland, and subjects provided written informed consent following the committee’s instructions. The study was conducted according to the principles of the Declaration of Helsinki. All potential subjects who donated BAT were screened for metabolic status, and only those with normal glucose tolerance and normal cardiovascular status (as assessed based on electrocardiograms and measured blood pressure) were included. The age range of the subjects was 23–49 years. We studied a group of 5 healthy volunteers (3 females, 2 males).

#### Human samples (RNAseq, refers to figures 6 and 7)

The acquisition of human data from cohorts 1 and 2 has been previously described [53]. Briefly, cohort 1 includes 318 individuals (249 female individuals, 69 male individuals; BMI range: 21.9 – 97.3 kg/m^2^, age range: 19-75 years) undergoing elective laparoscopic surgery during which subcutaneous (scWAT) and visceral, omental (visWAT) adipose depots were obtained. Cohort 2 and 3 are from the Leipzig Obesity Biobank. Cohort 2 includes 96 individuals (23 lean female individuals, mean age 43.3 ± 7.4 years, mean BMI 23.7 ± 1.3 kg/m^2^), 48 obese female individuals, mean age 42.9 ± 8.3 years, mean BMI 45.9 ± 6.1 kg/m^2^), 9 lean male individuals, mean age 44.7 ± 7.1 years, mean BMI 22.3 ± 1.8 kg/m^2^), and 16 obese male individuals, mean age 42.2 ± 7.6 years, mean BMI 44.9 ± 5.2 kg/m^2^)). Cohort 3 comprises 1415 individuals, with visWAT samples collected from 920 individuals (female lean (n = 45, mean age 62.1 ± 13.7 years, mean BMI 24.4 ± 2.9 kg/m^2^), female obese (n = 582, mean age 46.1 ± 11.8 years, mean BMI 48.5 ± 8.5 kg/m^2^), male lean (n = 44, mean age 61.4 ± 15.7 years, mean BMI 24.9 ± 3.5 kg/m^2^), and male obese (n = 249, mean age 48.2 ± 11.7 years, mean BMI 49.1 ± 8.5 kg/m^2^)) and scWAT samples taken from 814 individuals (female lean (n = 33, mean age 63.8 ± 12.9 years, mean BMI 24.7 ± 2.8 kg/m^2^), female obese (n = 545, mean age 46.7 ± 12.0 years, mean BMI 48.3 ± 9.2 kg/m^2^), male lean (n = 25, mean age 65.6 ± 14.3 years, mean BMI 25.5 ± 2.6 kg/m^2^), and male obese (n = 211, mean age 48.09 ± 12.4 years, mean BMI 49.5 ± 8.0 kg/m^2^)). Patients were classified as lean when their BMI was less than 30 kg/m^2^. For cohort 3, human single-end and rRNA-depleted RNA-seq data were prepared with a SMARTseq protocol [60, 61]. In brief, RNA was enriched and reverse transcribed by Oligo(dT) and TSO primers. cDNA was amplified by ISPCR primers and processed with Tn5 using Nextera DNA Flex kit. All libraries were sequenced on an Novaseq 6000 instrument at Functional Genomics Center Zurich (FGCZ). Approval for all 3 studies was obtained from the Ethics Committee of the University of Leipzig (approval no: 159-12-21052012) before the study and acquisition was performed in accordance with the declaration of Helsinki.

### Bioinformatic Analysis

#### Bioinformatic Analysis (RNAseq screening, refers to figure 1)

RNA quality was assessed using BioAnalyzer 2100 (Agilent); all samples had RIN values ≥ 8.5. 4 μg total RNA per sample were used for the TruSeq Stranded mRNA LT Sample Prep Kit (Illumina) to generate cDNA libraries according to the manufacturer’s protocol. Single read sequencing was carried out using Illumina/Solexa HiSeq 2000. High-throughput sequencing was conducted by the Biomedical Sequencing Facility, Vienna. RNAseq alignment, long non-coding quantification and differential expression analysis were performed as follows: Raw sequencing reads were aligned against the human hg38 genome using STAR aligner with default parameters [62]. The mapped reads were assigned to genes using featureCount from the Bioconductor package Rsubread [63]. All annotated lncRNAs were quantified across each condition, using hg38 annotation. Normalization and differential expression analysis were performed using the R/Bioconductor package DESeq2 [64]. Significance was assumed for an adjusted p-value < 0.01. Data of the hpAS treated with norepinephrine are publicly available sequencing data [36].

#### Bioinformatic Analysis (RNAseq human cohort 3, refers to figures 6 and 7)

Adapters and low quality bases of raw reads were trimmed using fastp v0.20.0 [65]. Only reads with a minimum read length of 18 nts and which surpass a quality cut-off of 20 were kept. The remaining reads were aligned against the human (GRCh38.p13) genome from GENCODE [66] applying the STAR alignment algorithm v2.7.4a [67], allowing 50 multiple alignments per read. Standard pre- and post-mapping quality control was computed using FASTQC v.0.11.4 (https://www.bioinformatics.babraham.ac.uk/projects/fastqc). Gene counts were conducted with featureCounts v2.0.1 (10.1093/bioinformatics/btt656) where multiple mapped reads were fractionally counted. Count data were homoscedastic normalized with respect to library size using the variance stabilizing transformation from DESeq2 v1.32.0 [64]. Correlation analyses were computed using the psych R package v2.1.6 with the spearman correlation coefficient and confidence intervals of 0.05. P-values were adjusted using the Holm’s method (http://www.jstor.org/stable/4615733). Visualization of correlation analysis was performed with the corrplot v0.90 and ggstatsplot v0.8.0 R packages under R version 4.1.

### Statistics

Data is presented as mean ± standard error of mean (s.e.m). Student’s t-test and 2-way ANOVA were used as indicated in the figure legends and performed using GraphPad Prism 9.0 and R 4.1.0. In vitro experiments were performed at least 3 times and pooled where possible. Differences were deemed significant with a P-value < 0.05 and indicated by an asterisk.

## Results

### Identification of AATBC as a human lncRNA linked to NST in vivo and in vitro

Investigating the phenotypic signatures of different adipose depots in humans as well as isolated adipocytes offers the opportunity to unravel novel lncRNAs implicated in the control of thermogenic plasticity. Therefore, we performed a combined RNAseq analysis of three different human models. We compared lncRNA expression profiles of (1) WAT and BAT ex vivo, (2) differentiated human primary adipose-derived stem (hpAS) cells treated with norepinephrine (NE) and (3) human adipose derived stem (hMADS) cells treated with forskolin (Figure 1A). To confirm the induction of the thermogenic phenotype, we quantified *UCP1* mRNA expression in all samples (Figure 1C). Other thermogenic markers such as citrate synthase (*CS*), cell death inducing DFFA like effector A (*CIDEA*), peroxisome proliferator-activated receptor gamma coactivator 1-α (*PPARGC1A*) and adipose markers such as peroxisome proliferator-activated receptor-γ (*PPARG*), perilipin 1 (*PLIN1*) and fatty acid binding protein 4 (*FABP4*) were quantified for quality control purposes (Supplementary figure G-X). After rigorous filtering (Supplementary figure 1A-D), we found 368 lncRNAs to be differentially regulated in WAT vs BAT, 105 lncRNAs in hpAS treated with NE compared to control cells, and 116 lncRNAs in hMADS stimulated with forskolin compared to control cells, respectively. Out of these differentially regulated set of lncRNAs, 25 candidates were regulated in the same fashion under all three conditions (Figure 1B). Among these, we found that the majority of lncRNAs had higher expression levels under thermogenic conditions (Figure 1F and Supplementary figure 1E,F). Interestingly, among the lncRNAs with unknown function in adipocytes, the lncRNA AATBC was one of the most prominent candidates, which was consistently upregulated in all three models of stimulated thermogenesis. Of note, we found that LNC00473 was consistently upregulated, which is in agreement with a previous study on human lncRNAs [47]. These findings indicate that our stratification strategy was valid. (Figure 1 F-L and Supplementary figure 1E,F). In summary, the thermogenic phenotype of adipocytes is characterized by an induction of lncRNA transcription. In the pool of upregulated lncRNAs, AATBC is consistently associated with increased thermogenesis in human adipose tissue and cultured adipocyte models.

**Figure 1:**
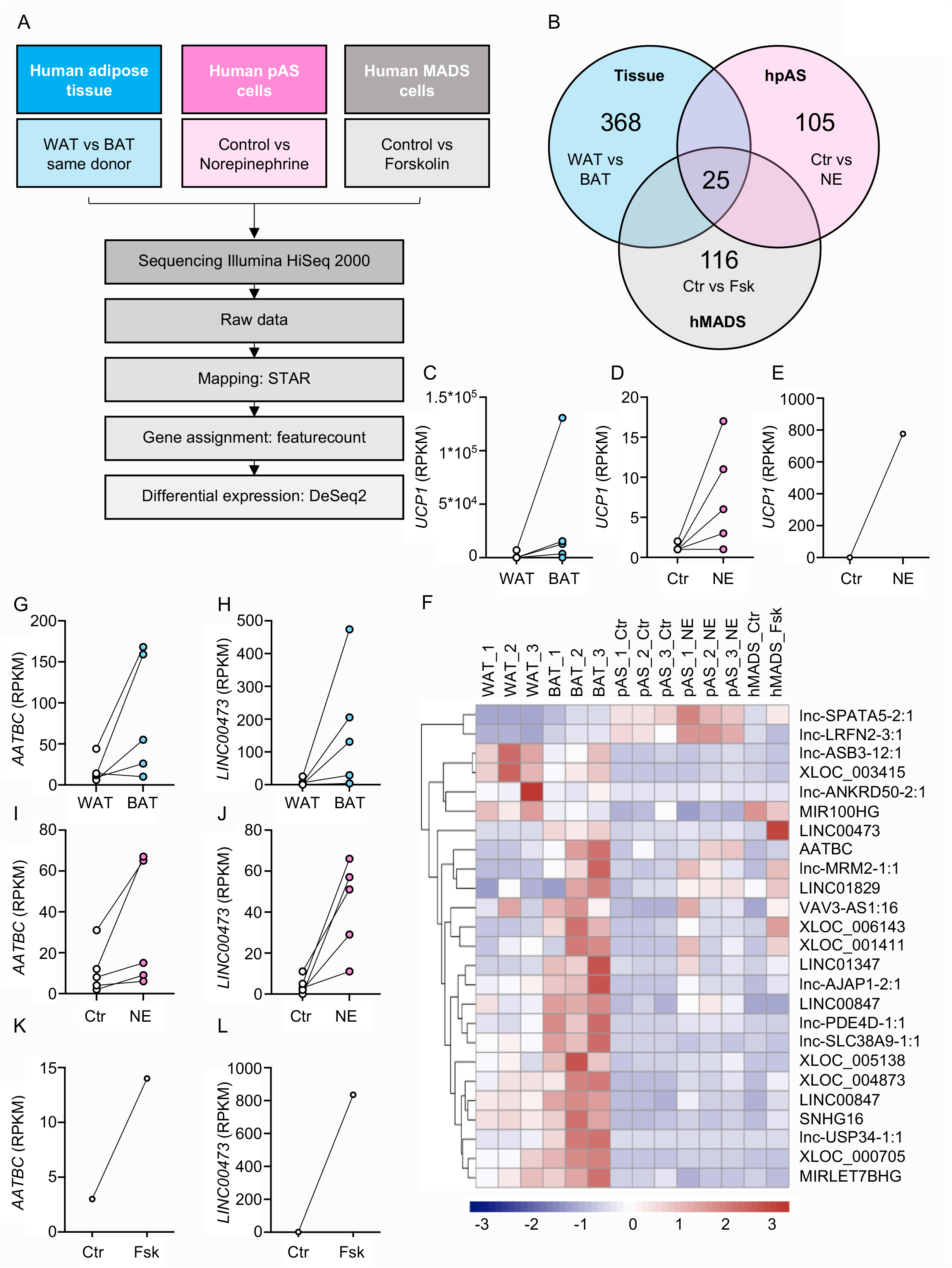
Identification of the lncRNA AATBC linked to thermogenesis in humans. **(A)** Experimental outline **(B)** Venn diagram of regulated lncRNA **(C-E)** *UCP1* expression in (C) human brown and white adipose tissue, (D) human primary adipocytes treated with 1 µM norepinephrine for 6 hours and (E) hMADS cells treated at day 15 of differentiation with 1 µM forskolin for 6 h (C: *n* = 5 patients, D: *n* = 5 replicates, E: *n* = 1 replicate) **(F)** Heatmap of the differential expression of lncRNAs (*n* = 1-3 replicates) **(G-L)** Expression of *AATBC* and *LINC00473* (G,H: *n* = 5 subjects; I,J: *n* = 5 replicates; K,L: *n* = 1 replicate).

### AATBC is a nuclear lncRNA induced by thermogenic activation and differentiation

As AATBC matched the candidate profile for a lncRNA associated with thermogenic features, we further explored the nature and regulation of AATBC expression in our in vitro models of human adipocytes. Confirming the results of the initial RNASeq screening, AATBC expression was higher in primary human brown adipocytes treated with NE (Figure 2A, B). As adipocyte browning is also induced by chronic PPARγ activation [68],we treated hSVF with the PPARγ agonist rosiglitazone during the differentiation (Supplementary figure 2A-C). Also, in this distinct model of adipocyte browning AATBC expression was higher under thermogenic conditions compared to controls cells differentiated to classical white adipocytes (Figure 2C). Using adipose tissue fractionation, we found that AATBC was expressed at higher levels in differentiated adipocytes compared to SVF cells (Figure 2D). Also, in hMADS cells we observed higher levels of AATBC in the respective models of thermogenic and regular differentiation (Figure 2E-F). *UCP1, PLIN1 and FABP4* expression levels served as validation markers for the thermogenic and regular differentiation, respectively (Supplementary figure 2D-K). Next to expression levels, the localization of lncRNAs is very important for understanding their function. To this end, we separated nuclear and cytoplasmic RNA from differentiated thermogenic hMADS cells (Figure 2J,K) and found that AATBC was enriched in the nuclear fraction (Figure 2I). The finding that AATBC is a nuclear lncRNA was corroborated by in situ hybridization of AATBC (RNAscope®) in the same hMADS cells. There, endogenous expression of AATBC was detected in the nucleus. (Figure 2L). This observation was confirmed by overexpression of AATBC, after which in most cells AATBC was found in the nucleus (Figure 2M). In summary, our data demonstrate that AATBC is a nuclear lncRNA and is induced by thermogenic activation and differentiation.

**Figure 2:**
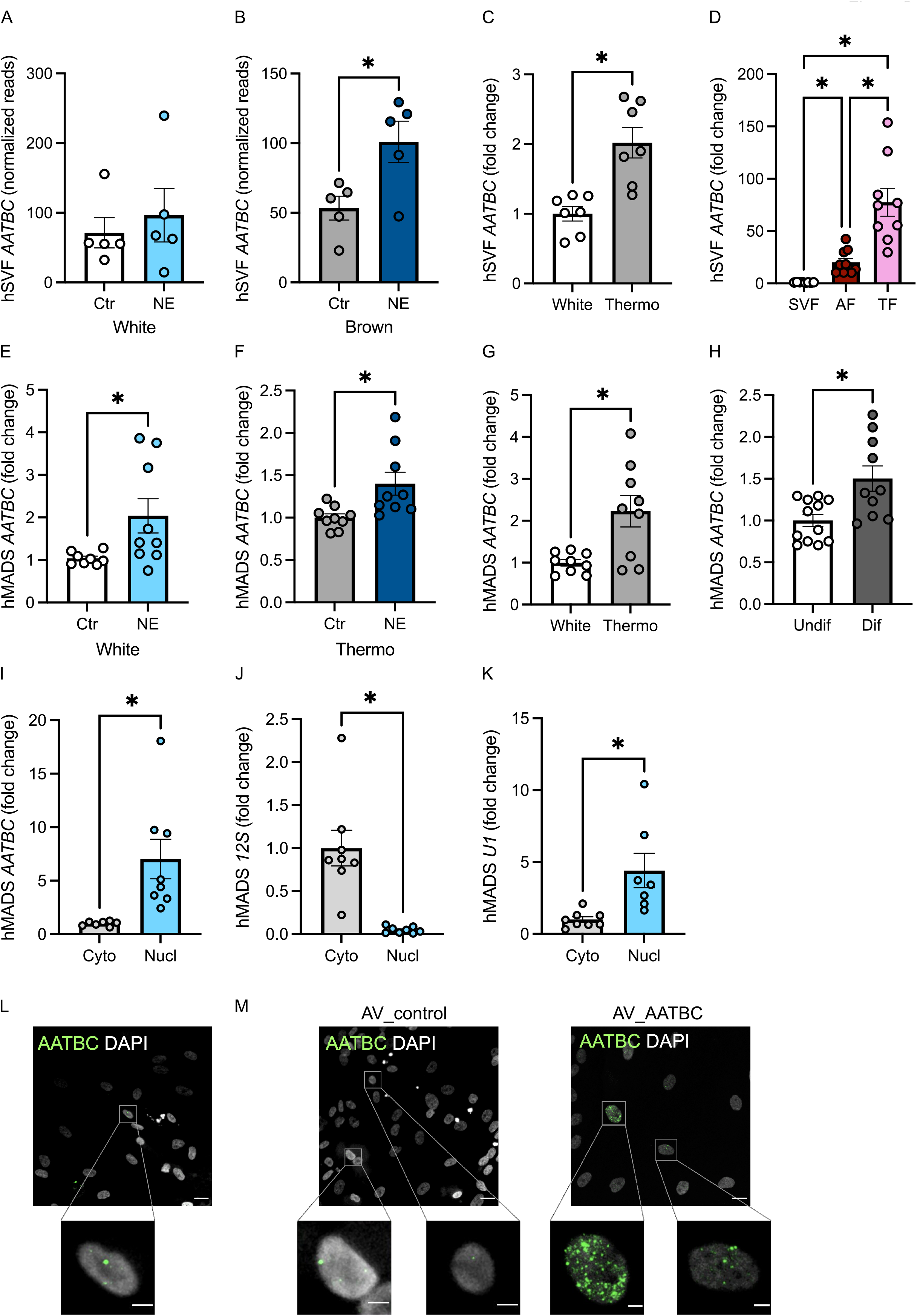
AATBC is associated with adipocyte browning and found in the nucleus. **(A, B)** *AATBC* expression in (A) white or (B) brown human stromal-vascular fraction (hSVF) cells treated with 1 µM norepinephrine for 6 h (*n* = 5 replicates) **(C)** *AATBC* expression in hSVF differentiated to a white and thermogenic phenotype (*n* = 7 replicates) **(D)** *AATBC* expression in the SVF, adipocyte fraction (AF) and total fraction (TF) of human white adipose tissue (*n* = 9 replicates) **(E**,**F)** *AATBC* expression in (E) white or (F) thermogenic human mesenchymal adipose stem (hMADS) cells treated with 1 µM norepinephrine for 6 h (*n* = 9 replicates) **(G)** *AATBC* expression in hMADS cells differentiated to a white and thermogenic phenotype (*n* = 9 replicates) **(H)** *AATBC* expression in undifferentiated and differentiated hMADS cells (*n* = 12 replicates) **(I-K)** Expression of *AATBC, 12S* and *U1* mRNA in the nucleus and cytosol of thermogenic hMADS cells (*n* = 8 replicates) **(L**,**M)** In situ hybridization (RNAScope®) of *AATBC* in thermogenic hMADS cells at (L) untreated conditions and (M) in thermogenic hMADS treated with adenovirus (AV)-mediated overexpression (OE) of *AATBC* (or control AV) (representative images from 3 independent experiments) (M). Scale bars: 20µm and 5µm for the magnified images. Statistical significance \**p* < 0.05 by 2-tailed, unpaired Student’s T-test **(A, C-K)** or 2-way ANOVA followed by Tukey’s test **(B)**.

### Modulation of AATBC expression regulates mitochondrial oxygen consumption

To investigate the biological significance of AATBC we performed loss and gain-of-function experiments in human adipocytes. We used anti-sense oligonucleotide (ASO)-mediated knockdown of AATBC in hMADS cells under thermogenic conditions, where otherwise AATBC expression is high, and adenoviral over-expression of AATBC in regular hMADS cells, where otherwise AATBC expression is low. (Figure 3A). Using two independent ASOs we found that AATBC silencing led to lower *UCP1* expression (Figure 3B, C). These observations were confirmed in primary cells derived from hSVF (Supplementary figure 3A,B). Similarly, an alternative approach using siRNA against AATBC led to similar results in hMADS cells (Supplementary figure 3E,F). As changes in UCP1 levels might indicate altered mitochondrial function, we assessed the bioenergetic profiles of hMADS with silenced AATBC and control cells. We found that silencing of AABTC diminished basal, isoproterenol-stimulated, uncoupled respiration as well as lower spare mitochondrial capacity (Figure 3D-H). Taken together, this data set indicates that loss of AATBC disrupts the thermogenic function of adipocytes. In the complementary approach, we found that after overexpression of AATBC *UCP1* expression remained unchanged (Figure 3I,J). Yet we observed higher basal respiration and higher spare mitochondrial capacity compared to control cells. However, isoproterenol-stimulated respiration and proton leak remained unchanged, which is in line with the unchanged UCP1 expression (Figure 3K-O). Of note, *PLIN1* and *FABP4* expression levels were largely unchanged, indicating that manipulation of AATBC did not interfere with adipogenesis per se (Supplementary figure 3C, D,G,H,I-L). Next, we investigated if the manipulation of AATBC function is linked to any transcriptional changes that could explain the mitochondrial phenotype. We silenced AATBC in thermogenic hMADS using siRNA and performed RNASeq. We found that despite a marked reduction in AATBC levels, the effect on global gene regulation was minor and only few genes were impacted by the loss of AATBC without any clear outcome (Supplementary figure 4A-C). We also performed a transcriptomic analysis of regular hMADS overexpressing AATBC and found lipid and triglyceride metabolism pathways to be regulated (Supplementary figure 4D-F). Regardless of these observations, the transcriptomic changes upon of AATBC manipulation were not immediately linked to mitochondrial function, potentially indicating the AATBC exerts its effects largely via posttranscriptional mechanisms.

**Figure 3:**
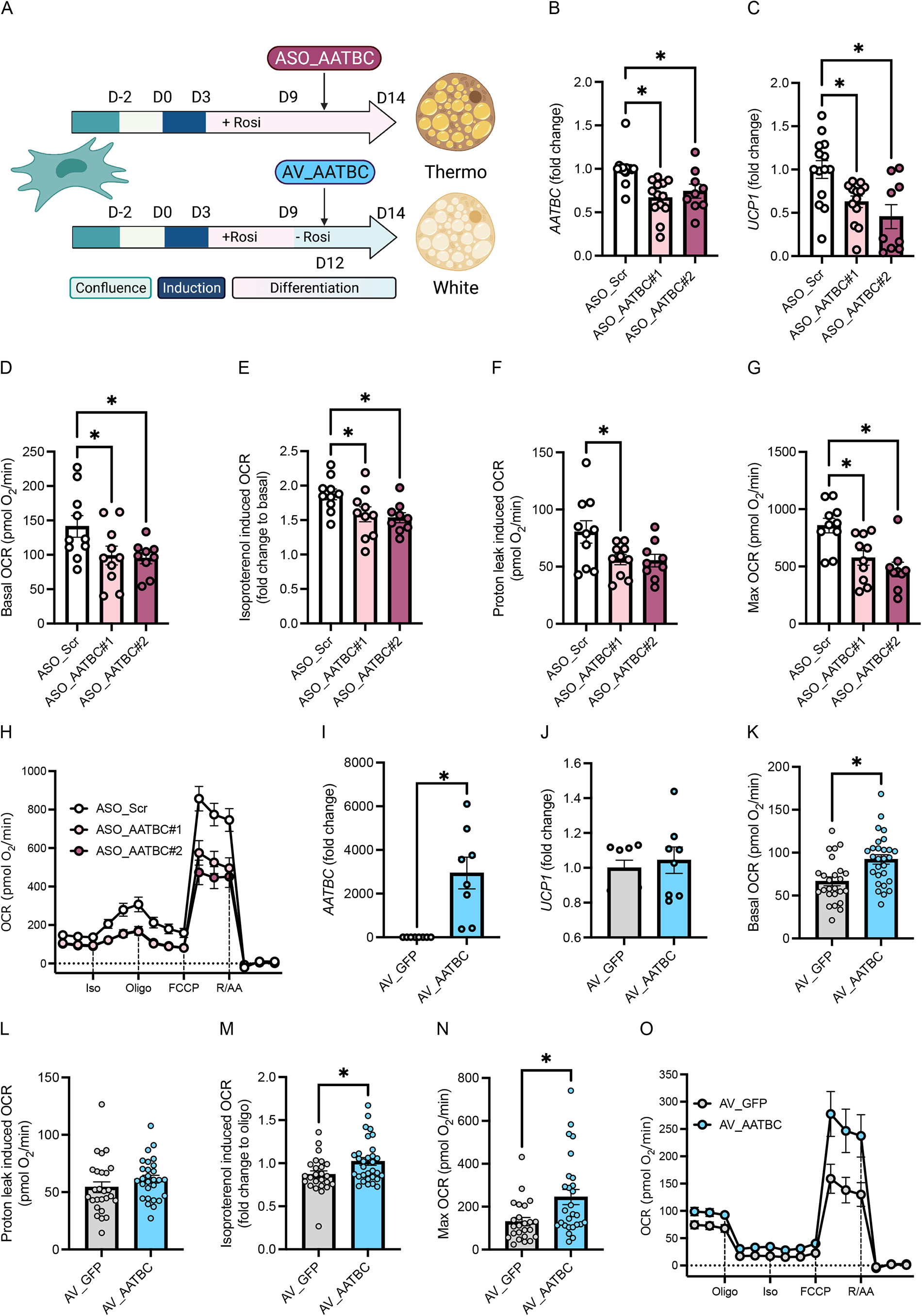
AATBC is a regulator of mitochondrial function. **(A)** Experimental outline **(B, C)** Expression of *AATBC* and *UCP1* during anti-sense oligonucleotide (ASO)-mediated knockdown of *AATBC* (*n* = 9-15 replicates) **(D-H)** Oxygen consumption rate (OCR) during ASO-mediated knockdown of *AATBC* visualized as (H) trace or cumulative at (D) baseline measurement, (E) isoproterenol-induced OCR, (F) oligomycin (Oligo) induced proton leak or (G) carbonyl cyanide-4-(trifluoromethoxy)phenylhydrazone (FCCP)-induced maximal respiration (D-G: *n* = 9 replicates, H: representative trace) **(I, J)** Expression of *AATBC* and *UCP1* during adenovirus (AV)-mediated overexpression of *AATBC* (*n* = 8 replicates) (**K-O)** OCR during ASO-mediated knockdown of *AATBC* visualized as (O) trace or cumulative at (K) baseline measurement, (L) isoproterenol-induced OCR, (M) OCR after oligomycin (Oligo) or (N) FCCP-induced maximal respiration (K-N: *n* = 27 replicates, O: representative trace) Statistical significance \**p* < 0.05 by 2-way ANOVA followed by Tukey’s test **(B-G)** or 2-tailed unpaired Student’s T-test **(I-N)**.

### Modulation of AATBC expression regulates mitochondrial dynamics

Considering that AATBC had only minor impact on the transcriptome of thermogenic adipocytes, we hypothesized that the mitochondrial phenotype is manifested at the posttranslational level. First, we quantified mitochondria DNA, but neither inhibition nor overexpression of AATBC led to altered mitochondrial DNA levels, suggesting that the number of mitochondria was unchanged under these conditions (Supplementary figure 5A,B). Next, we investigated mitochondrial dynamics, as mitochondrial fission and fusion events regulate mitochondrial respiration [22, 69]. Using TOM20 immunostaining, we found that silencing of AATBC in thermogenic hMADS was associated with higher numbers and length of mitochondrial branches as well as more mitochondrial junctions per cell area compared to control cells (Figure 4A-C). These observations illustrate a link between AATBC, mitochondrial network connectivity, and increased thermogenic function. As mitochondrial fusion is mediated in part by OPA1 and MFN2, we evaluated their protein levels. We found that compared to control cells the levels of both proteins were higher in hMADS cells upon knockdown of AATBC (Figure 4E-G), indicating a shift in mitochondrial dynamics towards fusion. We also measured protein levels of the different OXPHOS complexes, which remained unchanged except for complex V (ATP synthase), which was lower in hMADS cells with silencing of AATBC (Figure 4H, I). Complementary to this data set, overexpression of AATBC was associated with less mitochondrial connectivity (Figure 4J-M) and lower OPA1 but unchanged MFN2 and OXPHOS levels compared to control-infected cells (Figure 4N-R). In summary, these data demonstrate that modulation of AATBC expression regulates mitochondrial dynamics.

**Figure 4:**
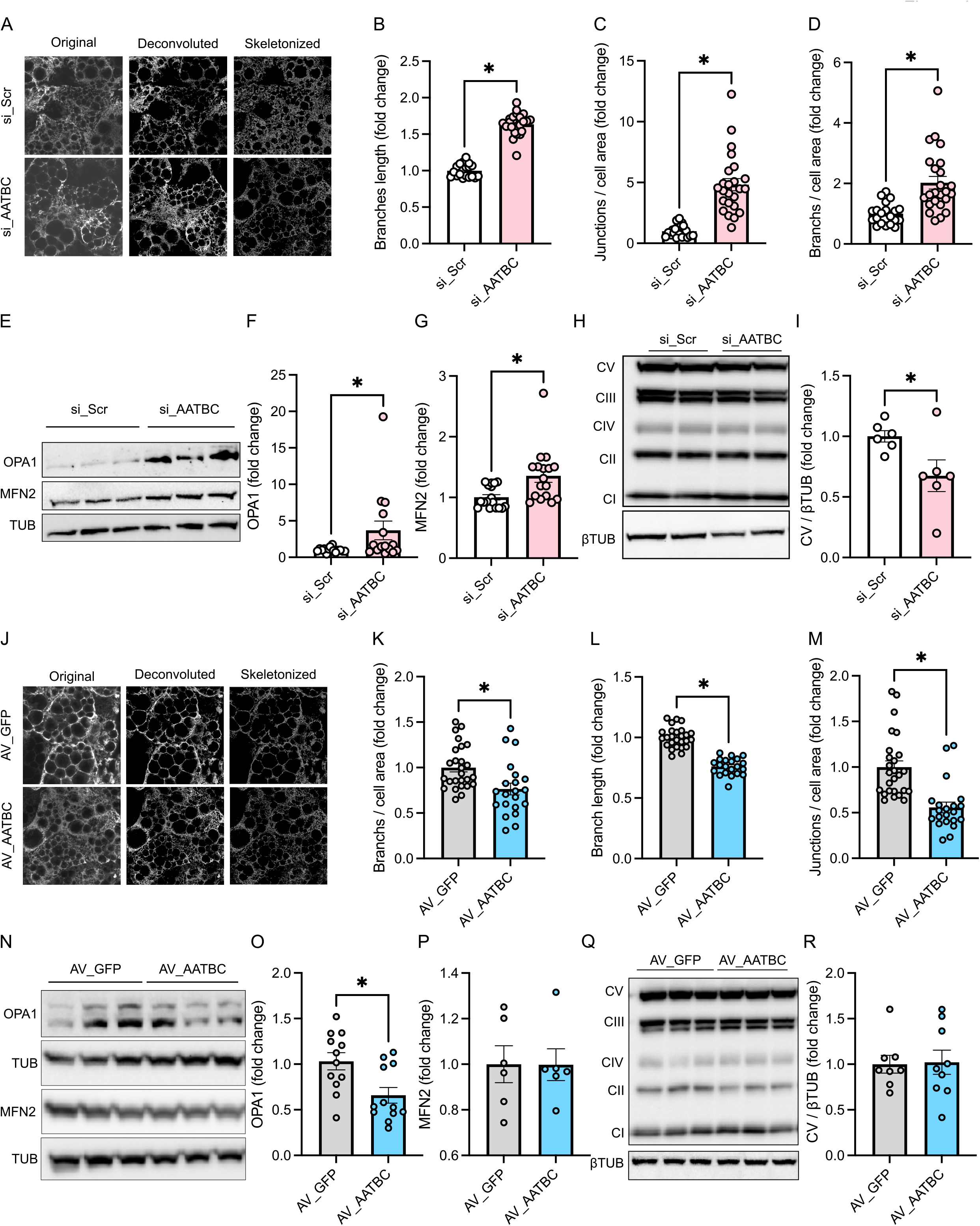
AATBC modulates mitochondrial dynamics by promoting fission. **(A)** TOMM20 staining and image analysis of white hMADS cells (representative image of 3 independent experiments) **(B-D)** Quantification of (B) length of mitochondrial branches, (C) junctions per cell area and (D) branches per cell area (*n* = 24 cells) **(E-G)** Immunoblot of OPA1, MFN2 and β-tubulin and the respective quantification normalized to β-Tubulin levels (E: representative immunoblot; F,G: *n* = 15 replicates) **(H**,**I)** Immunoblot of OXPHOS complexes and normalized to β-Tubulin levels (H: representative immunoblot; I: *n* = 6 replicates) **(J)** TOMM20 staining and image analysis of white hMADS cells (representative image of 3 independent experiments) **(K-M)** Quantification of (K) length of mitochondrial branches, (L) junctions per cell area and (M) branches per cell area (*n* = 24 observations) **(N-P)** Immunoblot of OPA1, MFN2 and β-Tubulin and the respective quantification normalized to β-Tubulin levels (N: representative immunoblot; O: *n* = 15 replicates; P: *n* = 6 replicates) **(Q**,**R)** Immunoblot of OXPHOS complexes and normalized to β-tubulin levels (Q: representative immunoblot; R: *n* = 6 replicates) Statistical significance \**p* < 0.05 by 2-tailed unpaired Student’s T-test **(B-D**,**F**,**G**,**I**,**K-M**,**O**,**P**,**R)**.

### Expressing human AATBC in mice impacts plasma leptin and triglyceride levels

AATBC is specific to humans, which complicates preclinical *in vivo* investigations in mouse models of metabolic disorders. Regardless, in a first attempt to overcome this obstacle we expressed AATBC in adipose tissue by surgically guided injection of adenovirus (AV) directly into interscapular BAT and inguinal subcutaneous (SCAT) adipose tissue of lean wild-type mice and subsequently performed several metabolic analyses (Figure 5A). In mice that received AATBC-AV we found marked, localized expression of AATBC lncRNA levels in both BAT and SCAT while in GWAT and liver only minor expression was detectable compared to control injected mice (Figure 5B). While body weight, glucose, and insulin tolerance as well as plasma insulin and adiponectin levels remained unchanged, we observed lower leptin levels (Figure 5C-I). In addition, in mice expressing AATBC we found higher plasma triglycerides while non-esterified fatty acids levels were unaltered (Figure 5J,K). However, the gene expression levels of *Ucp1* and adiponectin (*Adipoq*) were not affected (Figure 5L,M), indicating that the treatment has little to no deleterious effects. In line with the lower plasma leptin levels, also gene expression of leptin (*Lep*) was reduced in scWAT (Figure 5N). In conclusion, our proof-of-concept experiment shows AATBC expression in mouse adipose tissue influences systemic metabolism in vivo.

**Figure 5:**
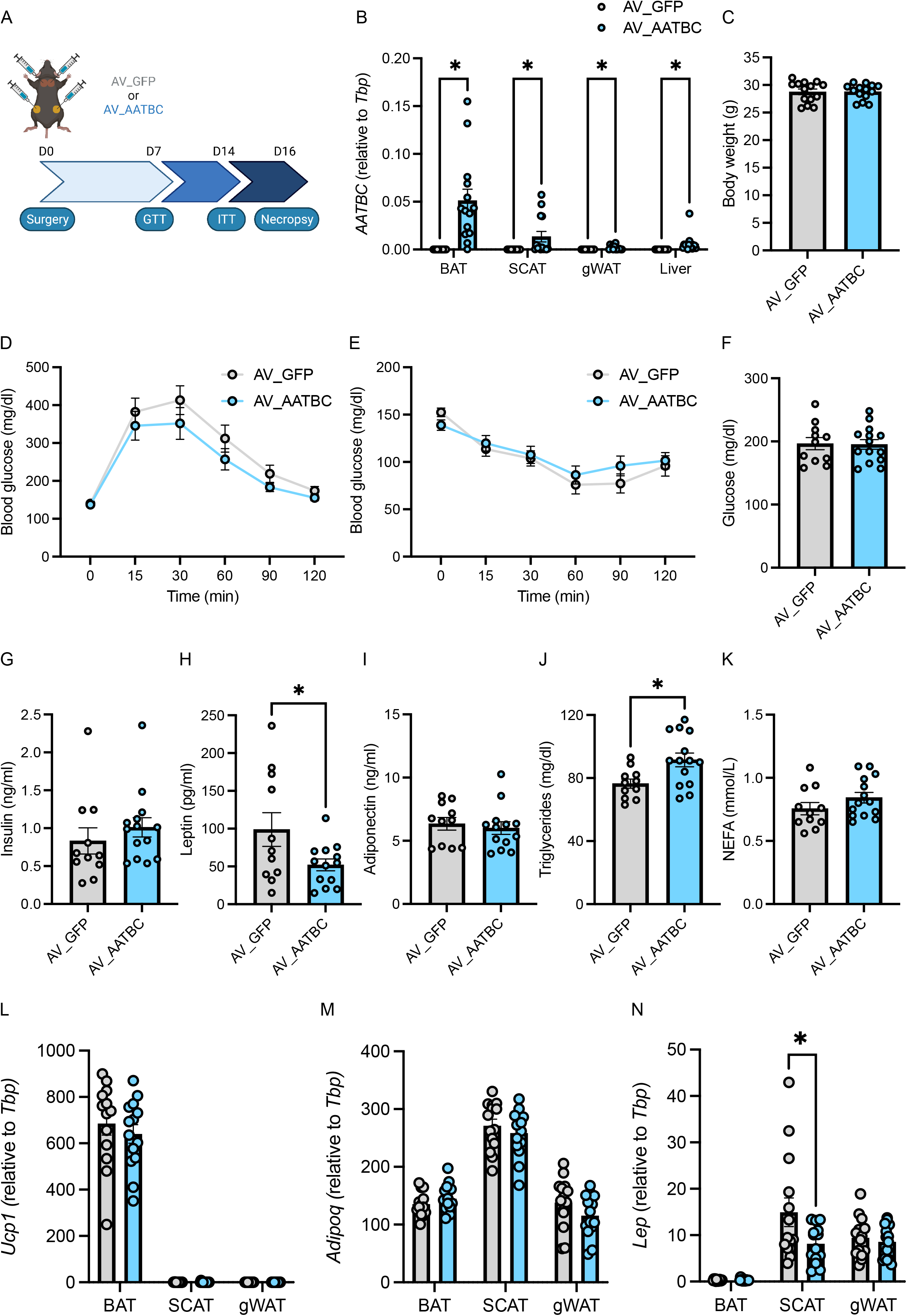
Expression of human AATBC in mouse adipose tissue impacts metabolism. **(A)** Experimental outline, **(B)** Expression of *AATBC* in adipose tissue depots and liver after injection of AV_GFP or AV_AATBC in brown adipose tissue (BAT) and subcutaneous adipose tissue (scWAT) **(C)** Body weight **(D)** Glucose tolerance test (GTT) **(E)** Insulin tolerance test (ITT) **(F-K)** plasma levels of metabolic parameters **(L-N)** *Ucp1, Adipoq* and *Lep* gene expression in adipose tissue (*n* = 12 animals per group) Statistical significance \**p* < 0.05 by 2-way ANOVA followed by Tukey’s test **(B**,**L-N)** or 2-tailed unpaired Student’s T-test **(C**,**F-K)**.

### AATBC is a novel obesity linked human lncRNA

To further corroborate the importance of adipocyte AATBC in systemic metabolism, we investigated the correlation of AATBC with different metabolic parameters in human obesity. Using cross sectional studies with subjects presenting a broad range of body mass index (BMI), we analyzed gene expression of AATBC by qPCR. In two independent cohorts we found that AATBC was expressed at higher levels in visceral (vis)WAT compared to scWAT (Figure 6A, B). Furthermore, the expression of AATBC was lower in visWAT of obese female subjects compared to lean controls (Figure 6B,D,E), an effect not found in scWAT (Figure 6C,F). In another separate cohort that included mainly obese subjects we were able to perform more in-depth analyses as in addition to anthropomorphic and plasma parameters, RNAseq data from adipose tissue samples was available. We found that AATBC was positively regulated with age and negatively regulated with body weight, body fat and the hip-waist ratio in visWAT of both females and males. Similar to our observation in mice, plasma levels of leptin were inversely correlated with AATBC, whereas adiponectin levels showed no correlation (Figure 6 G-J). In line with our previous observations, we found that AATBC was positively correlated with thermogenic markers such as *UCP1* and *PPARG1A* (Figure 7, A, B, D, E) and negatively correlated with the expression of genes coding for *ADIPOQ, LEP, FABP4* and *PPARG*. (Figure 7, C, F, G-L). These gene expression correlations were also found in scWAT but in the absence of correlations with the metabolic parameters (Supplementary figures 6 and 7). In summary, adipose AATBC is inversely linked to excess adipose tissue and leptin levels as well as correlates with thermogenic markers in humans.

**Figure 6:**
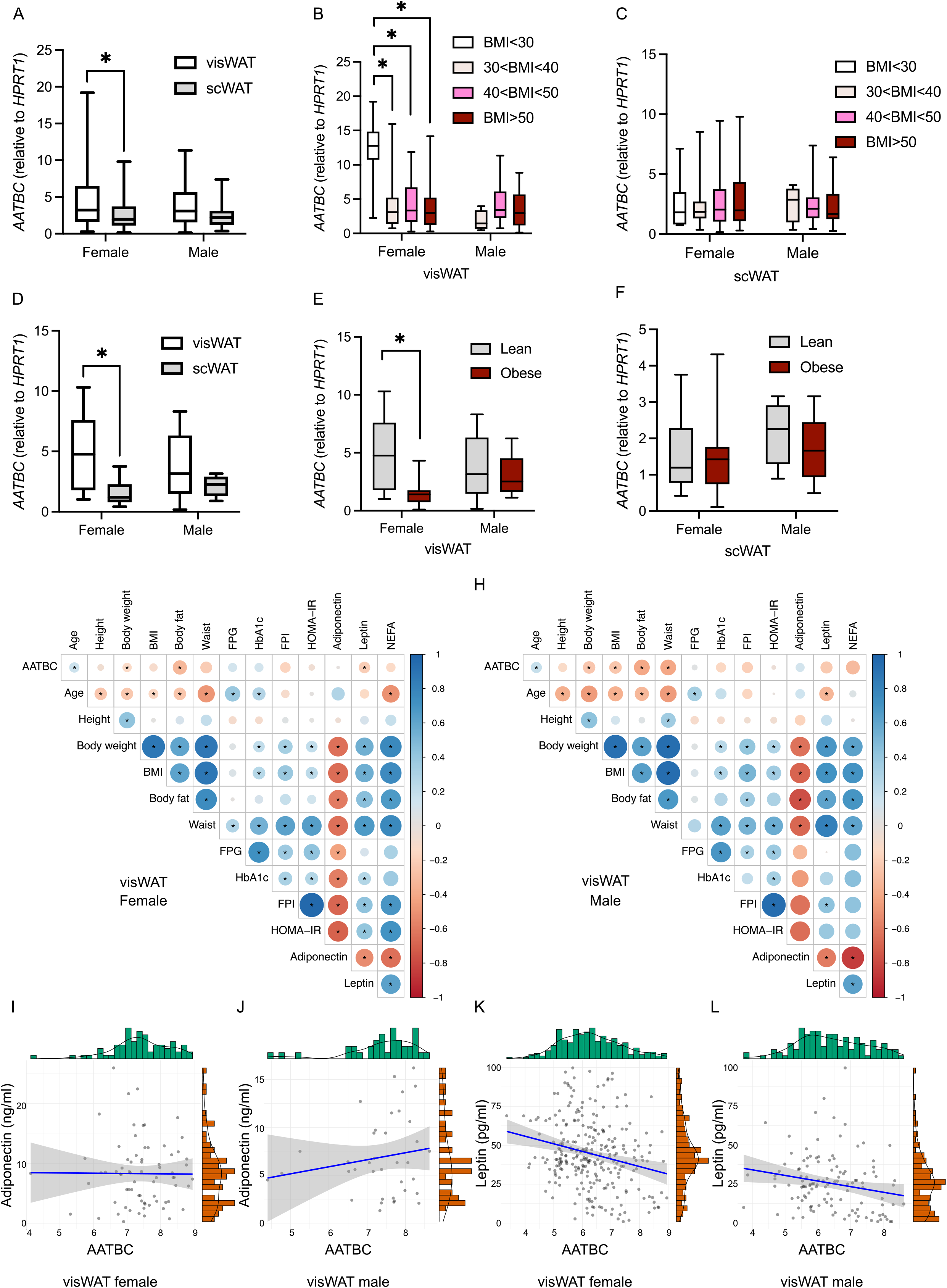
*AATBC* is linked to obesity and leptin levels in humans. **(A-C)** *AATBC* expression in visceral (visWAT) and subcutaneous (scWAT) white adipose tissue grouped by (A) sex (B,C) and body mass index (BMI) (*n* = 318) **(D-F)** *AATBC* expression in visWAT and scWAT of (A) male and female subjects (B,C) lean and obese subjects (*n* = 96 patients) **(G**,**H)** Correlation matrix of *AATBC* in visWAT of (G) female and (H) male subjects and metabolic parameters (females: *n* = 627, males: *n* = 293) **(I-L)** *AATBC* expression in visWAT of males and females correlated with (I,J) adiponectin and (K,L) leptin plasma levels (I: log_e_(*S*) = 11.02, *p* = 0.906, ρ_Spearman_ = 0.01, CI_95%_ [−0.22, 0.25], *n*_pairs_ = 72, J: log_e_(*S*) = 8.91, *p* = 0.465, ρ_Spearman_ = 0.12, CI_95%_ [−0.22, 0.44], *n*_pairs_ = 37, K: log_e_(*S*) = 15.16, *p* = 0.001, ρ_Spearman_ = −0.21, CI_95%_ [−0.32, −0.09], *n*_pairs_ = 267, L: log_e_(*S*) = 12.80, *p* = 0.011, ρ_Spearman_ = −0.23, CI_95%_ [−0.40, −0.05], *n*_pairs_ = 121) Statistical significance \**p* < 0.05 by 2-way ANOVA followed by Tukey’s test **(A-F)**.

**Figure 7:**
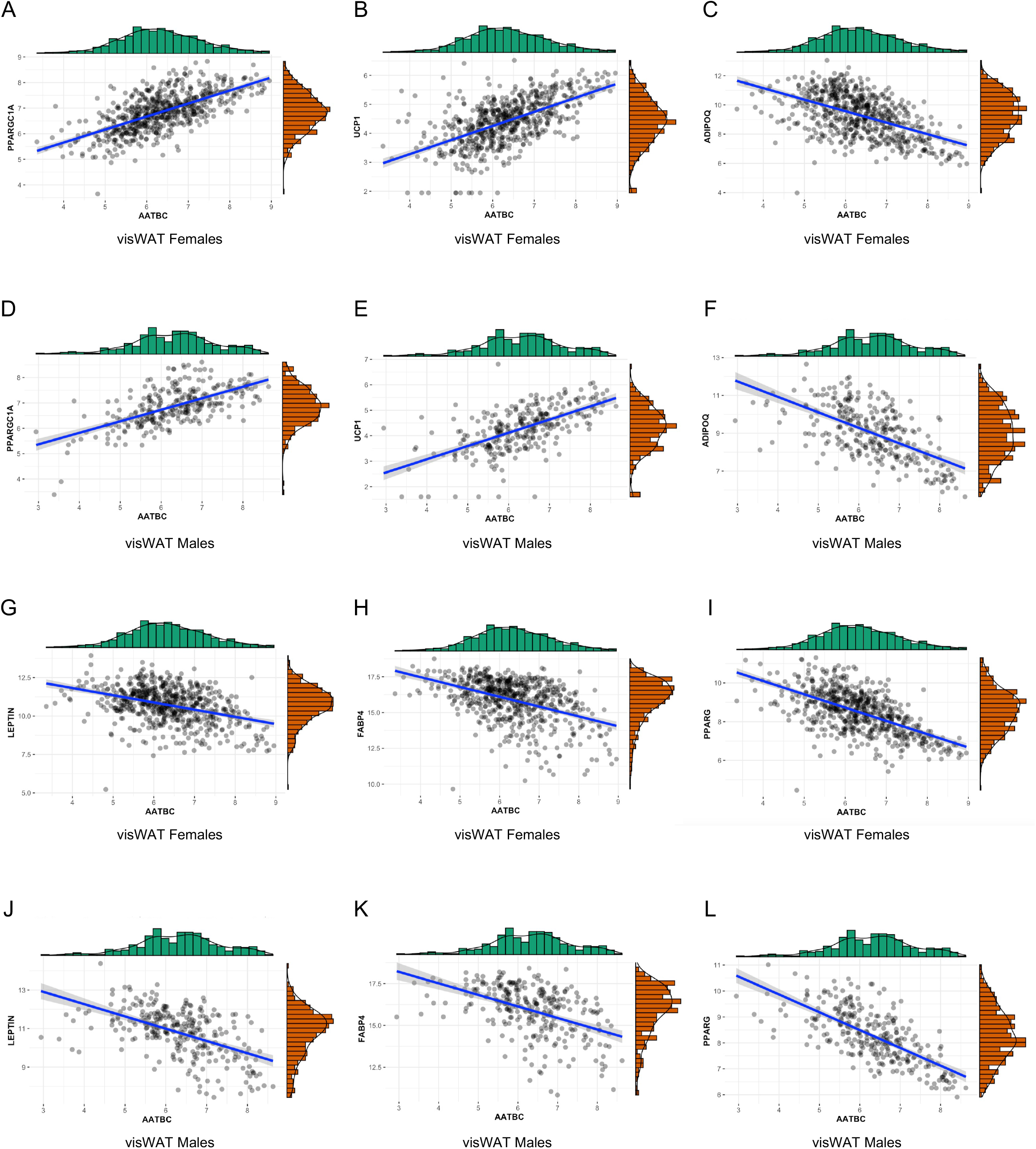
*AATBC* correlates with markers of metabolic health in visceral adipose tissue. **(A-L)** *AATBC* expression correlated with gene expression of the thermogenic markers *PPARGC1A* and *UCP1* and the obesity-related genes *ADIPOQ, FABP4, LEP* and *PPARG* in visceral adipose tissue of (A-C, G-I) females (*n* = 627) and (D-F, J-L) males (*n =* 293) (A: log_e_(*S*) = 16.47, *p* = 9.09e^−78^, ρ_Spearman_ = 0.65, CI_95%_ [0.61, 0.70]; B: log_e_(*S*) = 16.63, *p* = 1.37e^−60^, ρ_Spearman_ = 0.59, CI_95%_ [0.54, 0.64]; C: log_e_(*S*) = 17.94, *p* = 3.85e^−43^, ρ_Spearman_ = -0.51, CI_95%_ [-0.54, -0.45]; D: log_e_(*S*) = 14.30, *p* = 1.04e^−31^, ρ_Spearman_ = 0.61, CI_95%_ [0.53, 0.68]; E: log_e_(*S*) = 14.23, *p* = 7.55e^−35^, ρ_Spearman_ = 0.64, CI_95%_ [0.56, 0.70]; F: log_e_(*S*) = 15.71, *p* = 6.3e^−28^, ρ_Spearman_ = -0.58, CI_95%_ [-0.65, - 0.50]; G: log_e_(*S*) = 17.85, *p* = 4.95e^−22^, ρ_Spearman_ = -0.37, CI_95%_ [-0.44, -0.30]; H: log_e_(*S*) = 17.89, *p* = 6.46e^−29^, ρ_Spearman_ = -0.43, CI_95%_ [−0.49, −0.36]; I: log_e_(*S*) = 17.99, *p* = 7.06e^-60^, ρ_Spearman_ = −0.59, CI_95%_ [−0.64, −0.53]; J: log_e_(*S*) = 15.67, *p* = 1.1e^−21^, ρ_Spearman_ = −0.52, CI_95%_ [−0.60, −0.43]; K: log_e_(*S*) = 15.65, *p* = 1.81e^−19^, ρ_Spearman_ = −0.49, CI_95%_ [−0.58, −0.40]; L: log_e_(*S*) = 15.78, *p* = 2.37e^−43^, ρ_Spearman_ = −0.69, CI_95%_ [−0.75, −0.63]).

## Discussion

Adipocyte plasticity is a key component for the adaptation of metabolism to environmental cues. Thus, the functional features of the adipocyte are matched to the metabolic needs of the organism. This is especially prominent during the adaptation to cold, during which adipocytes adopt a more thermogenic phenotype, or, conversely, when temperature increases or other conditions of positive energy balance such as high-fat diet-induced obesity. If the energy balance is positive, whitening prevails over browning and the adipocytes undergo significant morphological and functional remodeling. Here, we show that the obesity-linked lncRNA AATBC is a regulator of adipocyte plasticity that enhances mitochondrial function in human adipocytes.

Mechanistic metabolic studies in humans are challenging and are further complicated if the gene of interest is absent in mice. To overcome these obstacles, we took advantage of hMADS cells which are a representative model of human adipocytes with the potential to switch between white and thermogenic phenotypes and confirmed our findings in primary cells isolated from human WAT and BAT. We demonstrate that in cultured human adipocytes, AATBC stimulates mitochondrial fission, oxygen consumption, and induces the expression of several markers of browning. Its nuclear localization suggests that AATBC exerts its effects by modulation of transcriptional events, but the outcomes of both loss and gain of AATBC function experiments on the global transcriptome were marginal and not linked to changes in genes that would explain the effects of AATBC on mitochondrial function [70]. Therefore, a non-transcriptional function for AATBC seems more likely. Indeed, lncRNAs can shuttle out of the nucleus and, for example, LNC00473 has been shown to physically present at the lipid droplet-mitochondrial interface [47]. However, we did not observe such a behavior for AATBC, as it remained strictly nuclear.

Likewise, lipolysis is a key feature of adipocyte function and we found that AATBC influences lipolysis both in vitro and in vivo. These functional effects were also mirrored in our adipocyte transcriptomic analysis of inhibition and overexpression of AATBC, potentially indicating that AATBC might have a dual role the regulation of adipocyte function. Lipolysis and mitochondrial function are two important pillars of adipocyte plasticity that are rapidly regulated but also change during chronic adaptations to cold or high-fat diet feeding [20, 71]. To investigate if AATBC is sufficient to alter adipocyte function in vivo, we expressed AATBC by a viral strategy in mice where AATBC is naturally absent. We found that AATBC expression altered plasma triglyceride and leptin levels, two important systemic parameters of adipocyte function. One could argue that expressing AATBC in mouse adipocyte might be artificial, as AATBC is a human lncRNA and could be non-functional in a mouse cell. However, it has been shown that expressing human lncRNAs in mice is useful for investigating lncRNA function [72] and we did not observe any adverse side effects in mice expressing AATBC. While more work is certainly needed to study the role of AATBC in non-shivering thermogenesis and/or obesity, ideally using a stable transgenic mouse model, our work here indicates that adipocyte AATBC plays role in systemic metabolism.

In humans, adipose AATBC expression is correlated with a thermogenic phenotype. This was also reflected in human primary adipocytes and hMADS cells. Regardless, AATBC is more expressed in visWAT compared to scWAT both in lean and obese patients, which implies that AATBC plays an important role in white adipocytes, too. In line with this notion, AATBC expression was correlated inversely with parameters of obesity and plasma leptin levels. Mitochondrial dynamics are equally important for proper white adipocyte function, as patients with mutations in *MFN2* display a fragmented mitochondrial network, excess adiposity, and paradoxical suppression of leptin expression [73]. In general, proper mitochondrial dynamics are critical for maintaining a healthy pool of mitochondria and efficient mitophagy [74]. To this end, it will be intriguing to study the role of AATBC in obesity-induced mitochondrial dysfunction, in adipocyte and potentially also other cell types as AATBC expression is not restricted to adipocytes. In conclusion, our study identifies AATBC as a novel lncRNA linked to adipocyte function and obesity with the potential to manipulate mitochondrial function and thermogenic plasticity in humans.

## Acknowledgements

We thank Ez-Zoubir Amri (Institut de Biologie Valrose, Université Nice Sophia Antipolis, France) for sharing hMADS cells. We thank Daniela Hass for outstanding technical support as well as Florian Pauler for valuable advice on the lincRNA RNAseq analysis. We thank the members of Bartelt Lab for enjoyable atmosphere and stimulating discussions. The figures were created using BioRender. MG was supported by an Alexander von Humboldt Foundation postdoctoral fellowship. P.F.P. was supported by the Deutsche Forschungsgemeinschaft (FI 1700/7-1, Heisenberg program). D.T. was supported by a grant from the German Research Association (TE912/2-2). S.H. was supported by the Helmholtz Future topic ‘Aging and Metabolic Programming, AMPro’, by the Deutsche Forschungsgemeinschaft Trans-Regio (TRR205) and the Deutsches Zentrum für Herz-Kreislauf-Forschung Standortprojekt Cardiometabolism. A.B. was supported by the Deutsche Forschungsgemeinschaft Sonderforschungsbereich 1123 (B10), the Deutsches Zentrum für Herz-Kreislauf-Forschung Junior Research Group Grant, and the European Research Council (ERC) Starting Grant PROTEOFIT. We apologize to colleagues whose work we could not cite due to space limitations.

## Conflicts of interest

The authors declare no conflicts of interest and that there are no relationships or activities that might bias, or be perceived to bias, their work.

## Author contribution statement

M.G., S.Ko., M.S., S.H. and A.B. conceptualized the study. M.G., S.Ko., and A.B. wrote the manuscript. M.G., S.Ko, and A.B. interpreted the work. M.G., S.Ko., Y.K., O.LT., M.G.L., A.H, M.K., J.H., S.Kh., D.T., P.F.P., W.S., AG, C.W., M.W., K.A.W., M.B., S.N., A.Z., C.G.C., M.S., S.H. and A.B. contributed to the acquisition and analysis of the data. All the authors revised the manuscript and approved the final version of the manuscript.

## Notes

### Competing Interest Statement

The authors have declared no competing interest.

