## Supplementary Figures for "The obesity-linked human lncRNA AATBC regulates adipocyte plasticity by stimulating mitochondrial dynamics and respiration"

Supplementary Figure 1

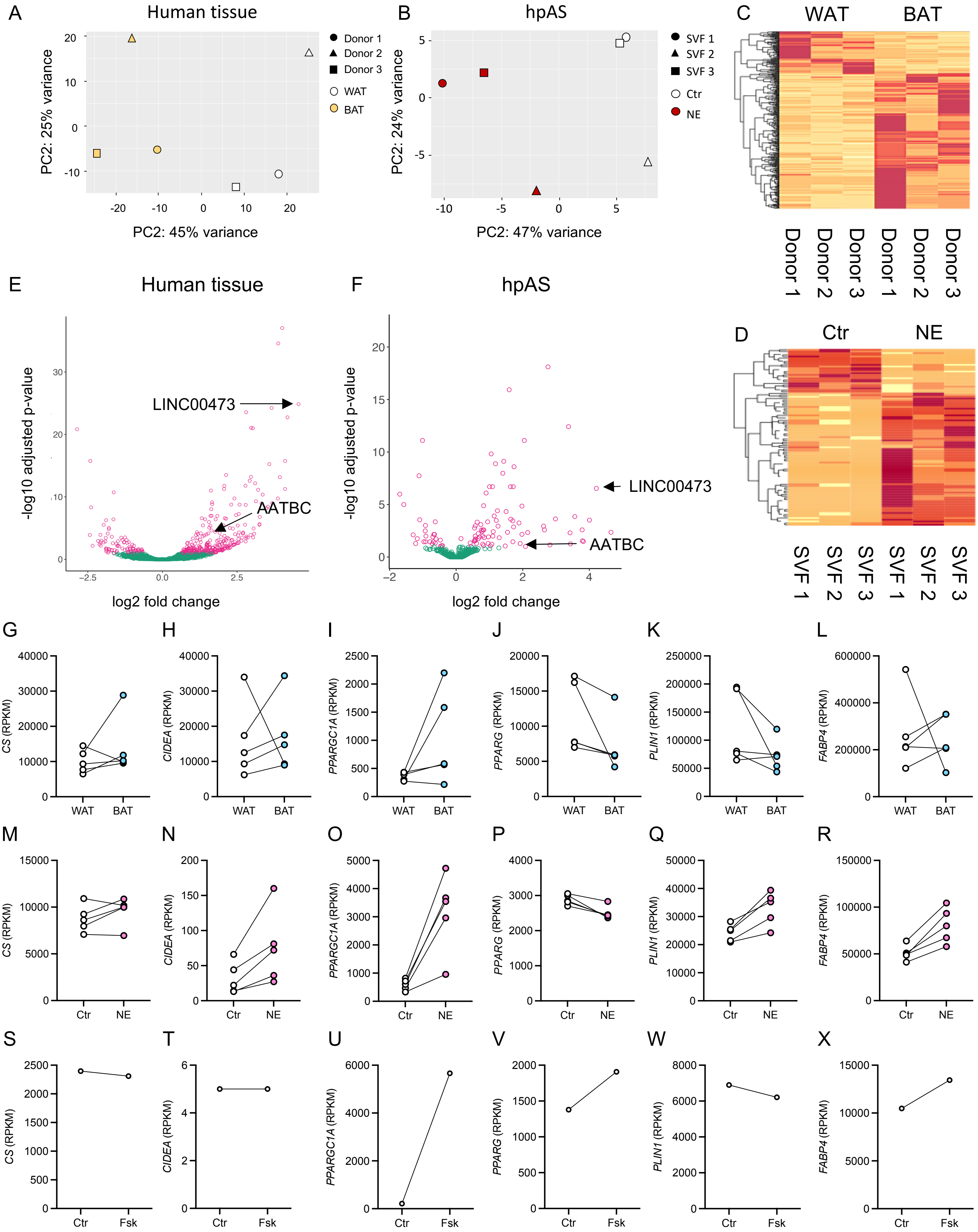

Supplementary Figure 1: Sequencing of lncRNA in different models of browning

**(A-F)** PCA, heatmap and volcano blot of lncRNA Sequencing of (A,C,E) human white and brown adipose tissue and (B,D,F) human primary adipocytes treated with norepinephrine **(G-X)** Gene expression of *CS, CIDEA, PPARGC1A, PPARG, PLIN1* and *FABP4* in (G-L) human brown and white adipose tissue (M-R) human primary adipocytes treated with 1 µM norepinephrine for 6 hours and (S-X) hMADS cells treated with 1 µM forskolin for 6 h.

Supplementary Figure 2

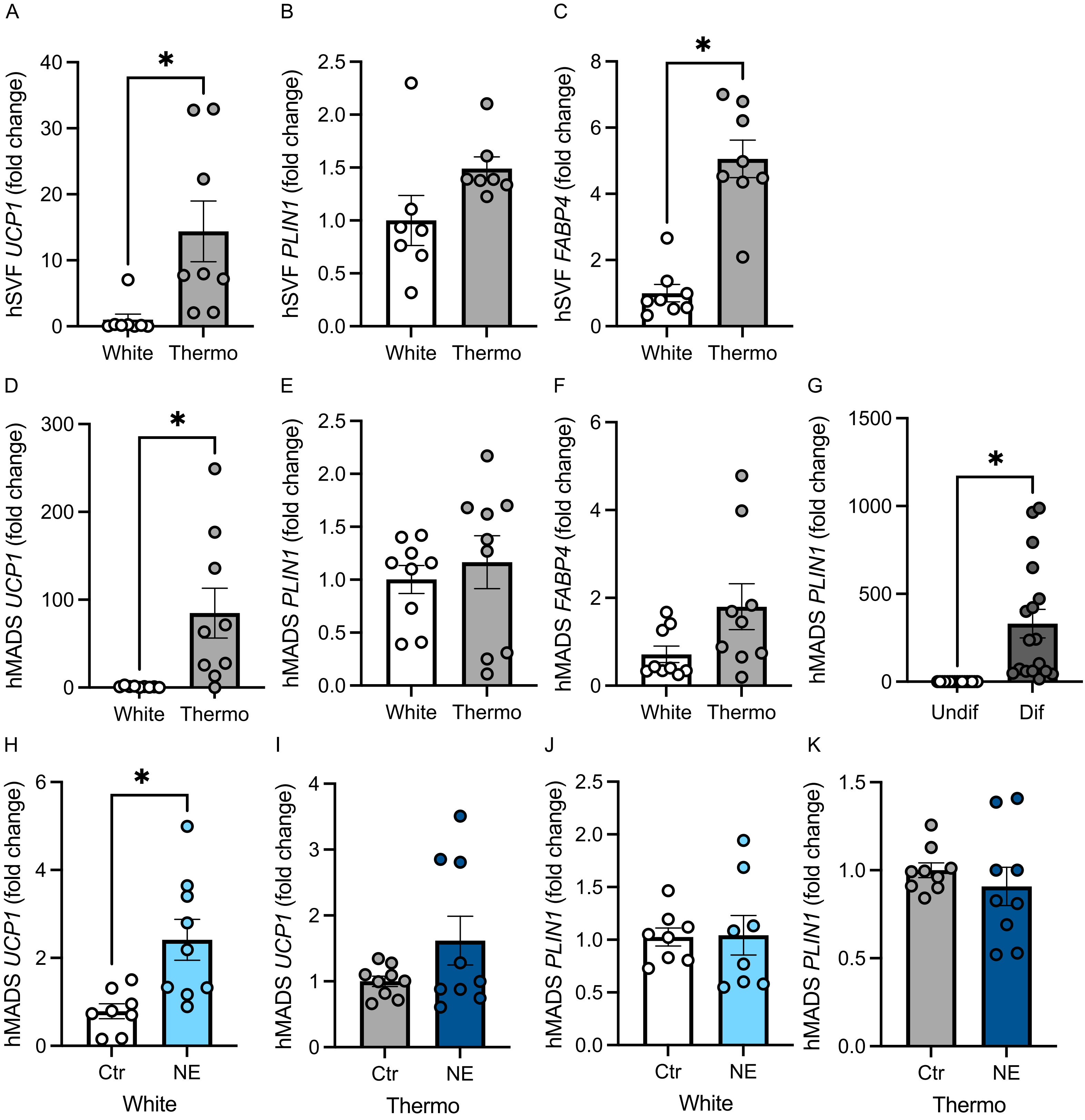

Supplementary Figure 2: Key markers of thermogenic differentiation in hSVF and hMADS cells
**(A-F)** Expression of *UCP1, PLIN1* and *FABP4* in (A-C) white and thermogenic hSVF and (D-F) white or thermogenic hMADS cells (A-C: $n$ = 8 replicates; D-F: $n$ = 9 replicates) **(G)** *PLIN1* expression in undifferentiated and differentiated hMADS cells ($n$ = 18 replicates) **(H-K)** Expression of *UCP1* and *PLIN1* in (H,J) white and (I,K) thermogenic hMADS cells treated with 1 μM norepinephrine for 6 h ($n$ = 9 replicates) Statistics: two-tailed unpaired $t$ test **(A,C,D,G,H)** Statistical significance is indicated by $^*p < 0.05$.

### Supplementary Figure 3

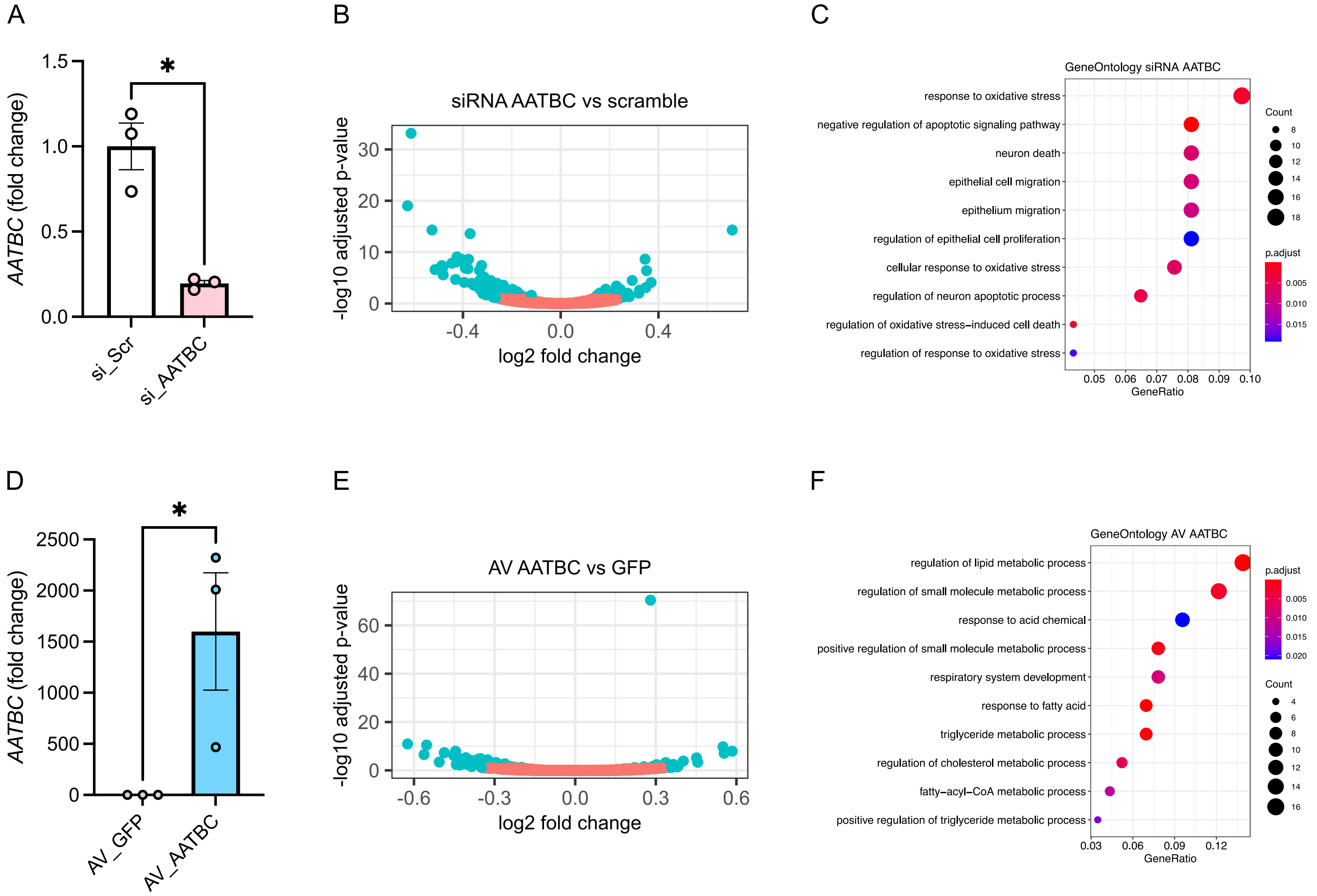

Supplementary Figure 3: mRNA Sequencing of knockdown and overexpression of *AATBC* in hMADS cells

**(A)** *AATBC* expression in hMADS cells upon siRNA-mediated knockdown of *AATBC* ($n$ = 3 replicates) **(B,C)** Vulcano blots and GO-Term analysis of knockdown of *AATBC* in hMADS cells (**D)** *AATBC* expression in hMADS cells upon AV-mediated over-expression of *AATBC* ($n$ = 3 replicates) **(E,F)** Vulcano blots and GO-Term analysis of over-expression of *AATBC* in hMADS cells Statistics: two-tailed unpaired $t$ test **(A,D)** Statistical significance is indicated by $*p < 0.05$.

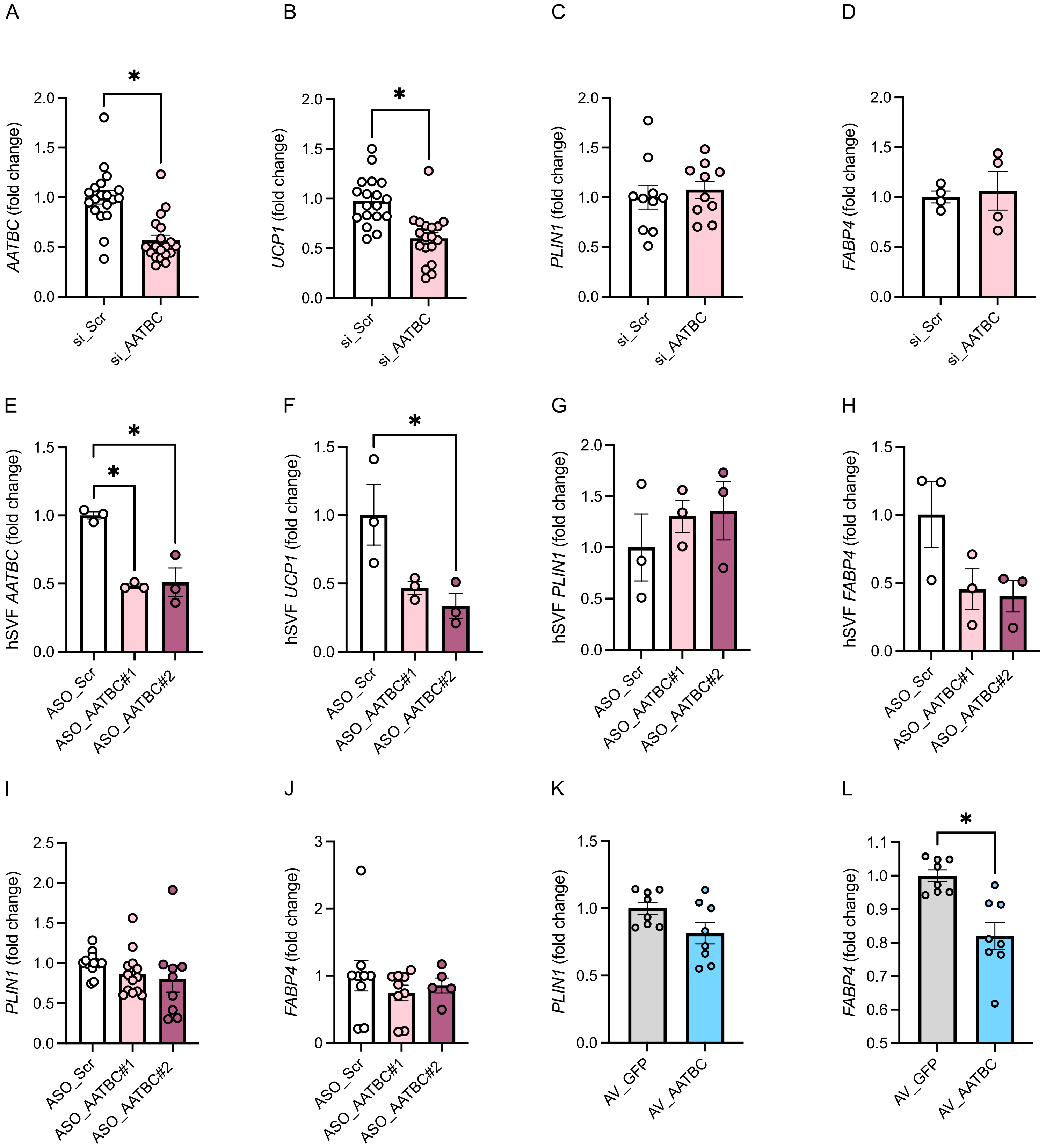
A
B
C
D
E
F
G
H
I
J
K
L
AATBC (fold change)
UCP1 (fold change)
PLIN1 (fold change)
FABP4 (fold change)
hSVF AATBC (fold change)
hSVF UCP1 (fold change)
hSVF PLIN1 (fold change)
hSVF FABP4 (fold change)
si_Scr
si_AATBC
ASO_Scr
ASO_AATBC#1
ASO_AATBC#2
AV_GFP
AV_AATBC
*

Supplementary Figure 4: Adipocyte gene markers upon modulation of *AATBC* gene expression **(A-H)** Gene expression of *AATBC, UCP1, PLIN1* and *FABP4* in (A-D) hMADS upon siRNA-mediated knockdown of *AATBC* and (E-H) in hSVF upon ASO-mediated knockdown of *AATBC* (A,B $n$ = 18 replicates; C $n$ = 10 replicates, D-H $n$ = 3 replicates) **(I-L)** *PLIN1* and *FABP4* expression in hMADS cells during (I,J) ASO-mediated knockdown and (K,L) AV-mediated over-expression of *AATBC* (I,J $n$ = 5-12 replicates; K,L $n$ = 8 replicates) Statistics: one-way-ANOVA with Tukey test **(A,B,E,F,L)** Statistical significance is indicated by $*p < 0.05$.

Supplemetary Figure 5

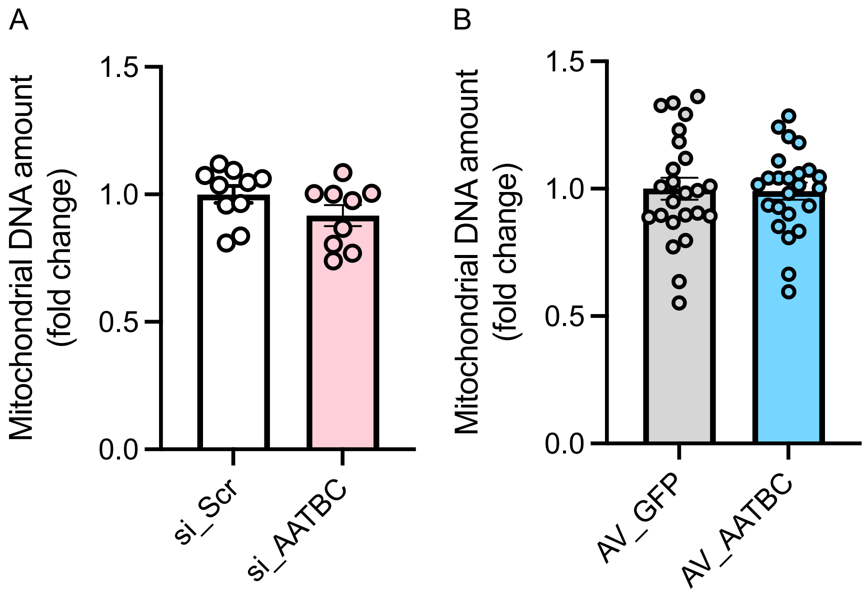

Supplementary Figure 5: Mitochondrial DNA content during modulation of *AATBC* expression

**(A,B)** Mitochondrial DNA amount during (A) siRNA-mediated knockdown and (B) AV-mediated over-expression of *AATBC* (A $n$ = 9 replicates; B $n$ = 24 replicates).

Supplementary Figure 6

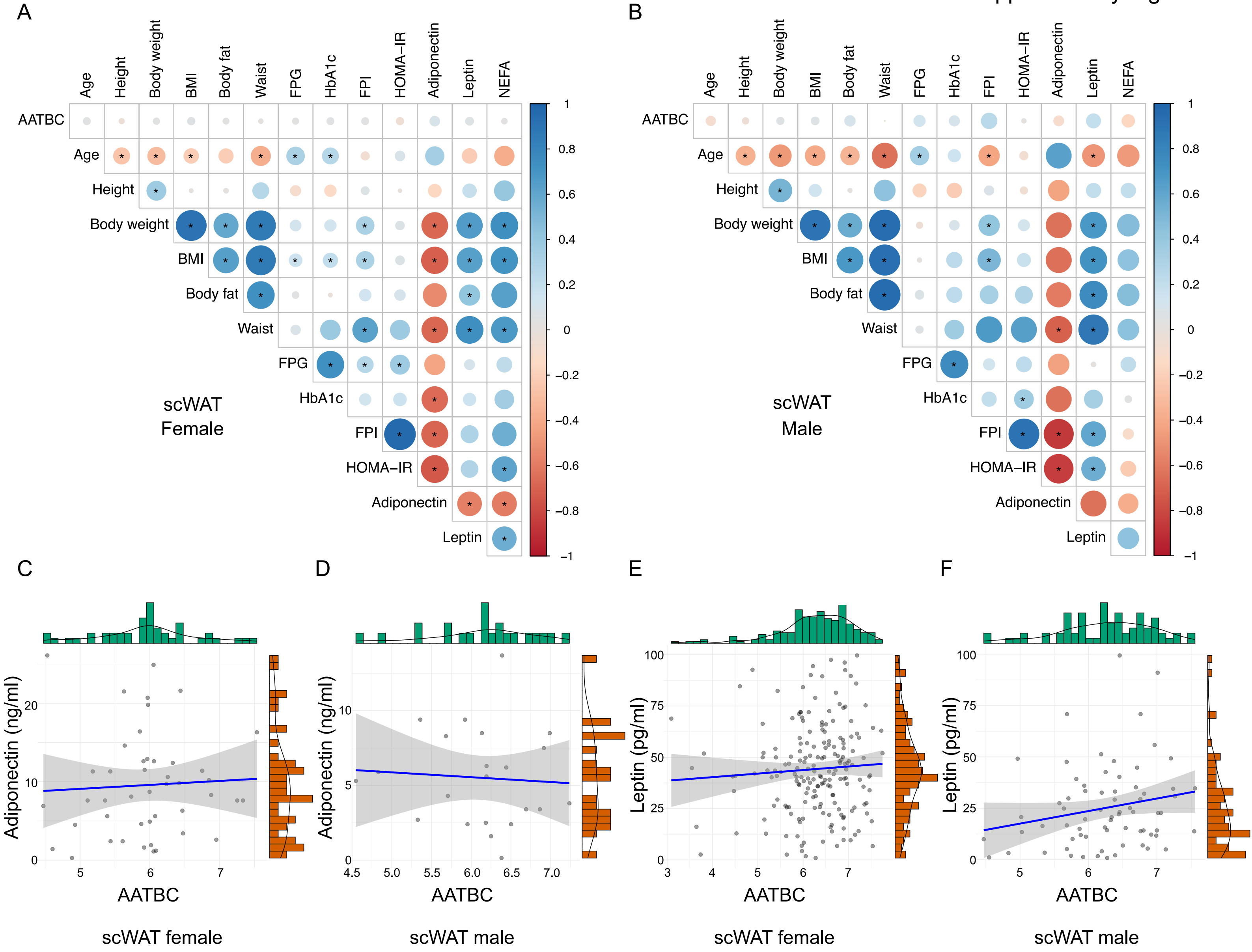

Supplementary Figure 6: Correlation of *AATBC* expression and metabolic parameters in human subcutaneous adipose tissue

**(A,B)** Correlation matrix of *AATBC* in subcutaneous adipose tissue of (A) female and (B) male patients and metabolic parameters (females: $n$ = 627 patients, males: $n$ = 293 patients, cohort 3) **(C-F)** Scatterplots of *AATBC* expression in subcutaneous adipose tissue of females and males correlated with (C,D) adiponectin and (E,F) leptin plasma levels (C: $\log_e(S) = 9.53$, $p = 0.530$, $\rho_{Spearman} = 0.10$, $CI_{95\%}$ [−0.21, 0.39], $n_{pairs} = 45$; D: $\log_e(S) = 7.58$, $p = 0.652$, $\rho_{Spearman} = -0.10$, $CI_{95\%}$ [−0.51, 0.35], $n_{pairs} = 22$, E: $\log_e(S) = 13.85$, $p = 0.377$, $\rho_{Spearman} = 0.06$, $CI_{95\%}$ [−0.08, 0.21], $n_{pairs} = 188$; F: $\log_e(S) = 10.73$, $p = 0.094$, $\rho_{Spearman} = 0.20$, $CI_{95\%}$ [−0.04, 0.42], $n_{pairs} = 70$).

Supplementary Figure 7

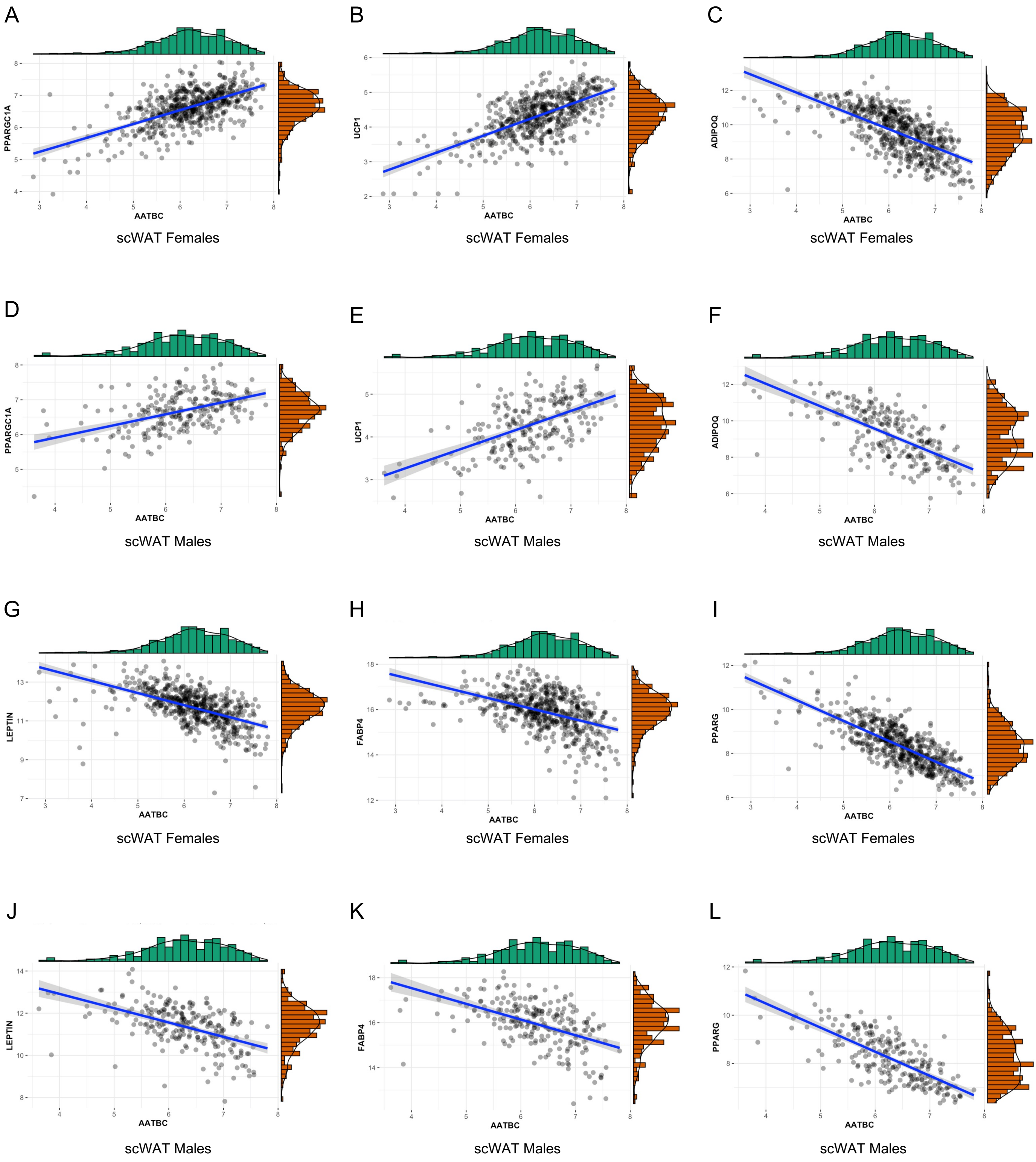

Figure 7: *AATBC* correlation with markers of metabolic healthy adipose tissue in human subcutaneous adipose tissue

**(A-L)** AATBC expression correlated with gene expression of the thermogenic markers PPARGC1A and UCP1 and the obesity related genes ADIPOQ, FABP4, LEPTIN and PPARG in subcutaneous adipose tissue of (A-C, G-I) females (n = 578) and (D-F, J-L) males (n = 236) (**A:** $\log_e(S) = 16.51$, $p = 7.4e^{-45}$, $\rho_{Spearman} = 0.54$, $CI_{95\%}$ [0.48, 0.60]; **B:** $\log_e(S) = 16.39$, $p = 1.17e^{-55}$, $\rho_{Spearman} = 0.59$, $CI_{95\%}$ [0.53, 0.64]; **C:** $\log_e(S) = 17.81$, $p = 1.17e^{-84}$, $\rho_{Spearman} = -0.69$, $CI_{95\%}$ [-0.73, -0.65]; **D:** $\log_e(S) = 14.00$, $p = 4.22e^{-13}$, $\rho_{Spearman} = 0.45$, $CI_{95\%}$ [0.34, 0.55]; **E:** $\log_e(S) = 13.85$, $p = 3.36e^{-18}$, $\rho_{Spearman} = 0.53$, $CI_{95\%}$ [0.42, 0.61]; **F:** $\log_e(S) = 15.12$, $p = 1.17e^{-33}$, $\rho_{Spearman} = -0.68$, $CI_{95\%}$ [-0.74, -0.60]; **G:** $\log_e(S) = 17.68$, $p = 8.44e^{-35}$, $\rho_{Spearman} = -0.48$, $CI_{95\%}$ [-0.54, -0.41]; **H:** $\log_e(S) = 17.68$, $p = 8.44e^{-35}$, $\rho_{Spearman} = -0.48$, $CI_{95\%}$ [−0.54, −0.41]; **I:** $\log_e(S) = 17.85$, $p = 3.29e^{-105}$, ρSpearman = −0.75, $CI_{95\%}$ [−0.78, −0.71]; **J:** $\log_e(S) = 15.03$, $p = 1.74e^{-19}$, $\rho_{Spearman} = -0.54$, $CI_{95\%}$ [-0.63, -0.44]; **K:** $\log_e(S) = 15.03$, $p = 1.74e^{-19}$, $\rho_{Spearman} = -0.54$, $CI_{95\%}$ [−0.63, −0.44]; **L:** $\log_e(S) = 15.15$, $p = 2.66e^{-40}$, $\rho_{Spearman} = -0.73$, $CI_{95\%}$ [−0.78, −0.66])
